# Integrated pangenome and population genomics reveal selection on standing genetic variation driving fiber flax-linseed divergence

**DOI:** 10.64898/2026.07.09.737549

**Authors:** Frank M. You, Chunfang Zheng, Tara Edwards, Pingchuan Li, Khalid Y. Rashid, Scott D. Duguid, Helen Booker, Sylvie Cloutier

**Affiliations:** Ottawa Research and Development Centre, Agriculture and Agri-Food Canada, Ottawa, ON, Canada; Morden Research and Development Centre, Agriculture and Agri-Food Canada, Morden, MB, Canada; Department of Plant Agriculture, University of Guelph, Guelph, ON, Canada

**Author notes:** Correspondence: Frank M. You; Sylvie Cloutier.

**Keywords:** flax, pangenome, population genomics, standing genetic variation, soft selective sweeps, morphotype divergence, telomere-to-telomere genome assembly, genome structural variation, quantitative trait nucleotide, genome-wide association study (GWAS), polygenic architecture

## Abstract

Flax (*Linum usitatissimum* L.) has been domesticated for dual end uses as linseed and fiber flax, yet the genomic basis of morphotype divergence remains unclear. Here, we constructed a morphotype-resolved pangenome by integrating three newly generated near telomere-to-telomere genome assemblies with 14 previously published ones. Despite substantial variation in assembly size, driven primarily by DNA transposons, gene content was highly conserved, with little evidence for significant morphotype-specific gene presence–absence variation. Population genomic analyses of 407 accessions revealed that fiber flax had reduced nucleotide diversity, extended linkage disequilibrium, and a more compact population structure relative to linseed, consistent with stronger selection and a narrower genetic base. Genome-wide differentiation was heterogeneous and concentrated in discrete regions. Integration of *F_ST_*, nucleotide diversity ratios, Tajima’s D, and genome-wide association signals identified morphotype-enriched genomic blocks distributed across the genome. Many candidate regions are primarily supported by directional shifts in nucleotide diversity rather than extreme differentiation, indicating selection on standing genetic variation. Genome-wide association analyses identified 1,712 unique quantitative trait nucleotides (QTNs), with predominantly small effect sizes and strong enrichment in gene-proximal regions, consistent with a polygenic architecture. Overall, fiber flax traits tend to be controlled by fewer loci with moderate-to-large effects, whereas linseed traits exhibit a more diffuse genetic architecture. Patterns of Tajima’s D further support non-classical selection dynamics, with predominantly positive values in linseed and localized negative values in fiber flax, consistent with selection on standing genetic variation. Together, our results suggest that flax morphotype divergence is driven primarily by selection on pre-existing allelic variation within a conserved gene repertoire. This study provides a comprehensive framework linking genome structure, population genomics, and trait architecture, and highlights the importance of standing genetic variation as a key resource for flax breeding and improvement.

## INTRODUCTION

Flax (*Linum usitatissimum* L.) is one of the earliest domesticated crops and has been cultivated for more than 10,000 years for two distinct end uses: linseed production and fiber extraction (Allaby et al., 2015; Zohary et al., 2012). Archaeological evidence indicates that flax was domesticated in the Fertile Crescent, initially selected for seed oil and subsequently diversified for multiple agricultural applications (Zohary et al., 2012). Pale flax (*Linum bienne* Mill.) has been identified as the wild progenitor of cultivated flax (Fu, 2011). Following domestication, flax diversified into the distinct morphotypes of the linseed and fiber types (Diederichsen and Ulrich, 2009; Soto-Cerda et al., 2013).

Fiber flax and linseed exhibit pronounced differences in plant architecture and physiology. Fiber flax cultivars are typically tall, slender, and unbranched with elongated internodes and long bast fibers, whereas linseed cultivars are shorter, highly branched, and optimized for oilseed production (Diederichsen and Ulrich, 2009; Soto-Cerda et al., 2013). These contrasting phenotypes suggest coordinated genetic changes affecting developmental regulation, cell wall biosynthesis, and metabolic pathways. However, the genomic basis underlying fiber flax–linseed divergence remains unclear.

Advances in long-read sequencing have enabled the generation of chromosome-scale and near telomere-to-telomere (T2T) genome assemblies, substantially improving resolution of complex genomic regions (Nurk et al., 2022; Rhie et al., 2021). In flax, several high-quality reference genomes are now available, enabling studies of genome structure and gene annotation (Lu et al., 2025; You et al., 2018; You et al., 2026). However, most analyses remain based on a single reference genome and therefore do not capture intraspecific variation such as structural variation (SV), presence–absence variation (PAV), and haplotype diversity, which are important for domestication and trait evolution (Alonge et al., 2020; Bayer et al., 2020).

Pangenome approaches provide a framework to characterize genomic diversity by integrating multiple assemblies to define conserved (core) and variable (accessory) gene sets (Golicz et al., 2016a; Tettelin et al., 2005). In several crop species, including rice, wheat, maize, and soybean, pangenome analyses have revealed substantial gene content variation associated with domestication and breeding (Bayer et al., 2020; Liu et al., 2020; Montenegro et al., 2017; Yang et al., 2025). Multiple chromosome-scale assemblies and preliminary pangenomes have recently been produced in flax, but the relative contributions of gene content variation and allele-frequency divergence to morphotype differentiation have not been investigated.

Addressing this question requires integrating pangenome analyses with population-scale genomic data. SV, PAV, and allele-frequency shifts may all contribute to phenotypic divergence, while demographic history and breeding practices may have shaped genetic diversity differently for each morphotype. For example, fiber flax breeding has often relied on a narrower genetic base and strong directional selection for stem-related traits, which may contribute to reduced genetic diversity and extended haplotype structure relative to linseed (Duk et al., 2021; Guo et al., 2019; Sertse et al., 2019; Zhang et al., 2020a).

In this study, we constructed a morphotype-resolved flax pangenome by integrating three newly generated near T2T genome assemblies representing the linseed cultivars Bison and Novelty, and the fiber flax cultivar Laura with 14 previously published assemblies. This pangenome represents 17 flax cultivars spanning fiber flax, linseed, and dual-purpose morphotypes. In addition, we analyzed a diversity panel of 407 flax accessions genotyped with approximately 1.7 million SNP markers to investigate population differentiation and signatures of selection between morphotypes.

This study addresses four key questions: (i) the conservation of gene content across flax cultivars; (ii) differences in gene presence–absence and structural variation between fiber flax and linseed morphotypes; (iii) differences in genetic diversity and population structure between morphotypes; and (iv) location of genomic regions showing signatures of differentiation associated with morphotype divergence. Together, these analyses provide a genomic framework for understanding flax diversification and for improving both fiber flax and linseed breeding.

## RESULTS

### *De novo* near telomere-to-telomere (T2T) genome assemblies of three flax cultivars

We generated *de novo* genome assemblies for two linseed cultivars (Bison and Novelty) and one fiber flax cultivar (Laura). PacBio HiFi sequencing produced 21.6–24.2 Gb of long-read data per cultivar, corresponding to ∼41–45× genome coverage, with mean read lengths of 9.5–11.5 kb, comparable to the reference genome CDC Bethune v3.0 (**Table S1**; **Fig. S1**).

All three assemblies resolved the expected 15 chromosomes with high contiguity and minimal unanchored scaffolds (**Fig. 1**; **Table S2**). Assembly sizes ranged from 489.26 Mb (Novelty) to 494.28 Mb (Laura), comparable to CDC Bethune v3.0 (504.00 Mb). Chromosome-scale pseudomolecules were represented by one to a few scaffolds and were supported by the genetic map of the Bison × Novelty recombinant inbred line population (**Fig. S2**; **Table S3**), indicating correct chromosome assemblies. The total length of anchored sequences (481.08–485.76 Mb) was also comparable to the reference (488.56 Mb). BUSCO assessment scores of 97.0–97.8% indicate near-complete representation of the gene space (**Fig. S3**).

**Fig. 1.**
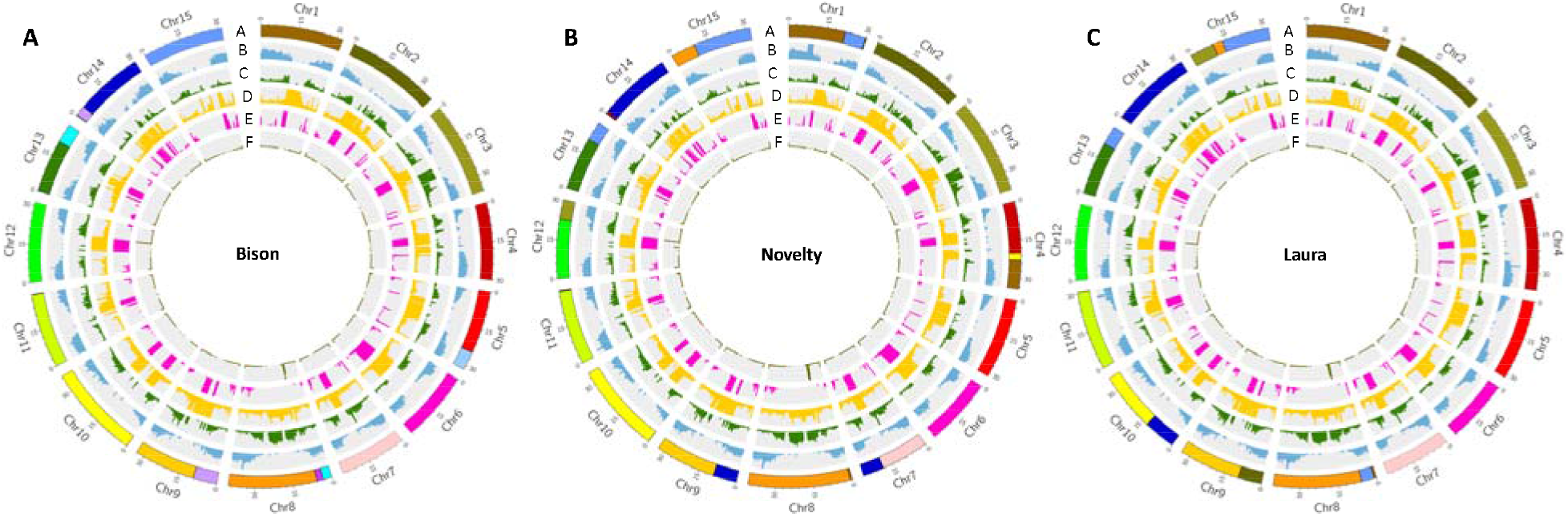
Circos plots illustrate near telomere-to-telomere (T2T) genome assemblies of three flax cultivars. **A:** linseed cultivar Bison. **B:** linseed cultivar Novelty. **C:** fiber flax cultivar Laura. Track A: chromosomes and assembled scaffolds; Track B: gene density; Track C: LTR retrotransposon density; Track D: DNA transposon density; Track E: dominant DNA transposon family *TE_00003234* (TIR/*hAT* superfamily); Track F: telomeric simple sequence repeat (SSR) motif distribution. Feature densities were calculated using a 100-kb sliding window.

### Assembly and chromosome size variation across the flax pangenome

To place these assemblies in a broader context, we integrated 14 previously published genomes with the three new assemblies to construct a pangenome of 17 chromosome-scale flax genomes spanning linseed, fiber, and dual-purpose flax cultivars (**Table 1**; **Table S4**). The relatively incomplete Heiya14 assembly was also included. Total assembly sizes ranged from 437.28 Mb (Ideo) to 504.00 Mb (CDC Bethune v3.0), with a median of 481.67 Mb. Chromosome-scale pseudomolecule sizes ranged from 386.65 Mb to 494.85 Mb (median 463.00 Mb). The three newly assembled genomes fell within this range, consistent with their high completeness.

**Table 1.** Summary of gene prediction and annotation for the flax pangenome, incorporating 17 diverse cultivars.

| Cultivar | Morphotype | Origin | Assembly<br>size (Mb) | Chr<br>size<br>(Mb) | Total<br>genes | HC<br>genes | LC<br>genes | Total gene<br>transcripts | Annotated<br>gene<br>transcripts | Annotated<br>(%) | Genes<br>with<br>GO | GO % | Genes<br>with<br>KO | KO<br>% |
| --- | --- | --- | --- | --- | --- | --- | --- | --- | --- | --- | --- | --- | --- | --- |
| CDC<br>Bethune<br>v3.0 | Linseed | Canada | 504.00 | 488.55 | 67,152 | 33,515 | 33,637 | 71,777 | 54,130 | 75.41 | 28,094 | 51.90 | 26,024 | 48.08 |
| **Bison** | Linseed | Canada | 490.06 | 481.94 | 65,732 | 33,418 | 32,314 | 70,299 | 52,678 | 74.93 | 27,949 | 53.06 | 25,891 | 49.15 |
| **Novelty** | Linseed | Canada | 489.26 | 481.08 | 65,386 | 33,397 | 31,989 | 69,969 | 52,836 | 75.51 | 28,067 | 53.12 | 25,962 | 49.14 |
| T397 | Linseed | India | 494.85 | 494.85 | 62,636 | 32,268 | 30,368 | 67,140 | 50,378 | 75.03 | 26,458 | 52.52 | 24,462 | 48.56 |
| K-1531 | Linseed | Russia | 412.43 | 373.81 | 56,512 | 33,449 | 23,063 | 61,139 | 49,969 | 81.73 | 26,950 | 53.93 | 24,895 | 49.82 |
| Line3896 | Linseed | Russia | 434.23 | 386.65 | 60,023 | 34,042 | 25,981 | 64,649 | 52,236 | 80.80 | 27,195 | 52.06 | 25,527 | 48.87 |
| Neiya9 | Linseed | China | 474.08 | 463.00 | 60,695 | 34,932 | 25,763 | 65,633 | 52,915 | 80.62 | 28,225 | 53.34 | 26,129 | 49.38 |
| Attila | Linseed | France | 441.56 | 417.26 | 60,505 | 33,349 | 27,156 | 65,126 | 51,809 | 79.55 | 27,145 | 52.39 | 25,068 | 48.39 |
| Marquise | Linseed | France | 441.45 | 420.72 | 60,312 | 33,292 | 27,020 | 64,858 | 50,646 | 78.09 | 27,231 | 53.77 | 25,180 | 49.72 |
| K-3018 | Dual | Yugoslavia | 489.13 | 487.19 | 65,228 | 33,421 | 31,807 | 69,820 | 52,987 | 75.89 | 28,263 | 53.34 | 26,127 | 49.31 |
| Gaosi | Dual | China | 482.50 | 482.50 | 65,228 | 33,305 | 31,923 | 69,865 | 53,084 | 75.98 | 28,234 | 53.19 | 26,156 | 49.27 |
| **Laura** | Fiber | Netherlands | 494.28 | 485.76 | 66,019 | 33,356 | 32,663 | 70,566 | 52,953 | 75.04 | 28,195 | 53.25 | 26,125 | 49.34 |
| Atlant v2.0 | Fiber | Russia | 491.33 | 491.13 | 65,797 | 33,375 | 32,422 | 70,566 | 53,730 | 76.14 | 28,166 | 52.42 | 26,106 | 48.59 |
| YY5 v2.0 | Fiber | China | 454.96 | 423.10 | 68,658 | 33,696 | 34,962 | 73,304 | 55,645 | 75.91 | 28,685 | 51.55 | 26,874 | 48.30 |
| Heiya14 | Fiber | China | 303.82 | 291.28 | 44,226 | 32,951 | 11,275 | 48,648 | 48,402 | 99.49 | 26,653 | 55.07 | 24,428 | 50.47 |
| Bolchoi | Fiber | France | 452.84 | 422.41 | 60,894 | 21,680 | 39,214 | 65,470 | 51,735 | 79.02 | 27,772 | 53.68 | 25,693 | 49.66 |
| Ideo | Fiber | France | 437.28 | 408.09 | 58,750 | 32,202 | 26,548 | 63,249 | 49,935 | 78.95 | 26,926 | 53.92 | 25,068 | 50.20 |

3 Chr, chromosome; GO, gene ontology; KO, KEGG Ortholog; HC: high-confidence; LC: low-confidence; Dual: dual-purpose flax; Fiber: 4 fiber flax

Most chromosomes were similar in size across cultivars except for Heiya14, although moderate variation was observed for some chromosomes (**Fig. S4**). For example, chromosome 3 ranged from 19.91 Mb to 42.83 Mb and chromosome 4 from 17.72 Mb to 39.91 Mb, whereas the size of chromosomes 1, 8, and 9 were similar across all genotypes. Overall, the new assemblies are consistent with the range of structural variations observed across flax genomes.

### Gene content and composition of the flax pangenome

Gene prediction for the three newly assembled genomes identified ∼65,000–66,000 genes, of which ∼33,400 were classified as high-confidence (HC) genes, comparable to CDC Bethune v3.0 (∼67,000 total genes; ∼33,500 HC genes) (**Table 1**). Genes were enriched toward chromosome arms, with lower densities in centromeric and pericentromeric regions (**Fig. 1**).

Across the pangenome, total gene numbers varied from ∼56,000 to ∼67,000), whereas HC gene sets were consistent (∼32,000–34,000 per cultivar) (**Table 1**). Functional annotation assigned putative functions to ∼74–82% of transcripts, with approximately half associated with gene ontology (GO) terms or KEGG orthologs, indicating consistent annotation quality across genomes.

Pangenome analysis of 17 cultivated flax genomes identified 71,992 orthogroups, with 33,910–41,478 orthogroups per cultivar (**Fig. 2**; **Table S5**). Orthogroups were classified into five pangenome categories based on their frequency of occurrence across genomes: core (present in all genomes), soft-core (present in 15/17 genomes), shell (3-14/17), rare (2/17), and private (unique to a single genome). The core genome comprised 30,792 orthogroups (∼52.8%), which, together with soft-core orthogroups accounted for ∼71.2% of all gene families (**Fig. 2A; Table S5**).

**Fig. 2.**
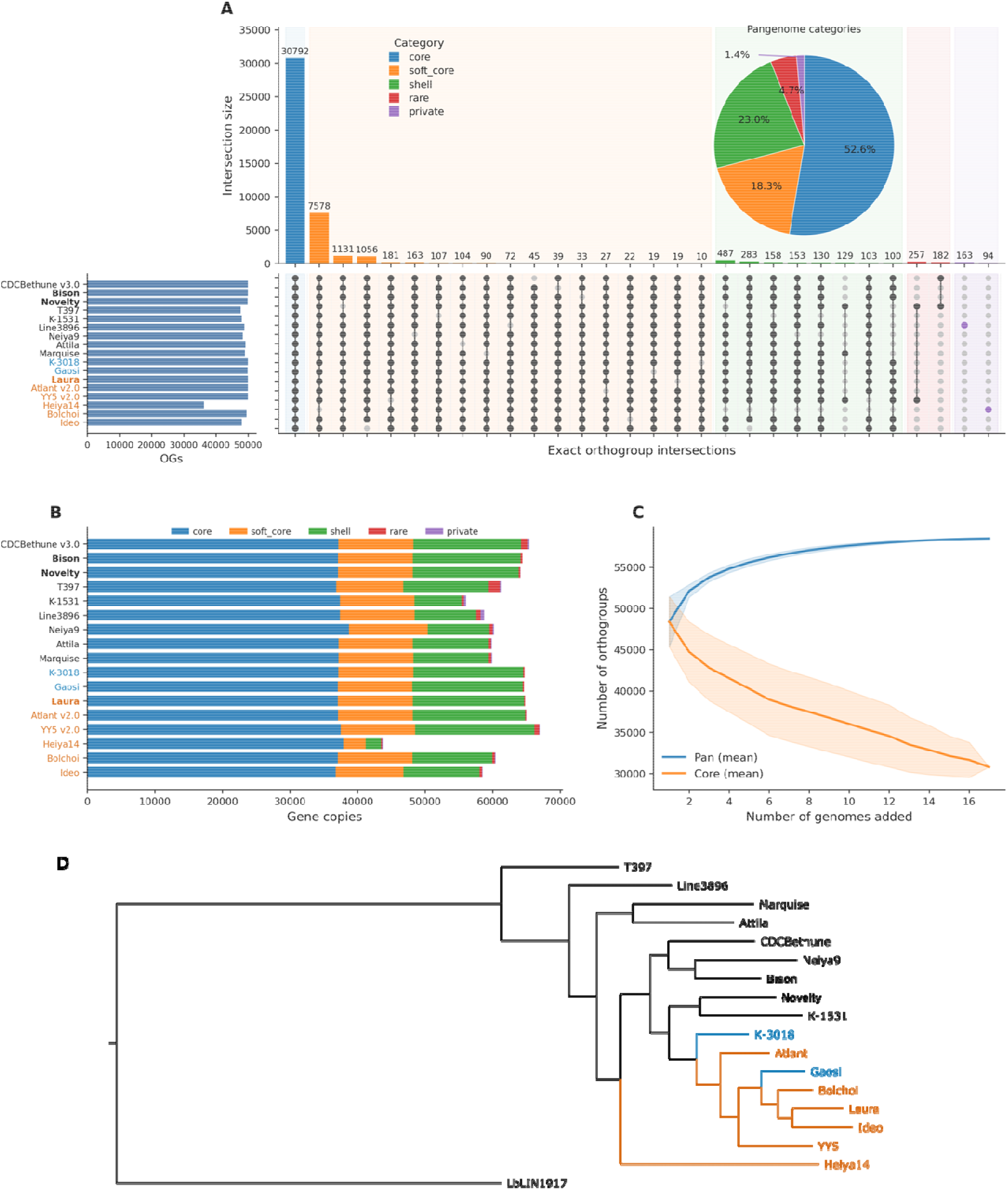
Orthogroup composition and pangenome structure of flax cultivars. **A:** Orthogroup intersections across 17 flax genomes visualized using an UpSet plot. The top histogram shows the number of shared orthogroups as designated by the matrix below which indicates the genomes included in each intersection. The pie chart summarizes the overall proportion of orthogroups belonging to each category in the flax pangenome. **B:** Gene copy composition across genomes according to pangenome categories. **C:** Pangenome accumulation curves based on sequential genome addition. Shaded regions represent variation among random genome addition permutations. **D:** Maximum likelihood phylogenetic tree of 17 cultivated flax (*L. usitatissimum*) cultivars and one pale flax (*L. bienne*) accession LIN1917 used as the outgroup (LbLIN1917). The tree was constructed using RAxML based on protein sequences from 20,178 single-copy genes under the LG+G+F substitution model. Branch support was assessed with 1,000 bootstrap replicates, and all nodes received 100% bootstrap support. Cultivar names are shown in black for linseed, blue for dual and orange for fiber flax.

Variation in soft-core gene content was associated with a subset of genomes showing orthogroup absence. Heiya14, T397, and Ideo lacked genes in 7,578, 1,131, and 1,056 soft-core orthogroups, respectively (**Fig. 2A**). In Heiya14, the 7,937 genes missing relative to CDC Bethune v3.0 were distributed across all chromosomes (324–726 genes per chromosome; mean ∼528). In contrast, missing genes in T397 and Ideo were concentrated in specific regions. In T397, 878 of 1,148 genes (76.48%) from 1,131 orthogroups were associated with an ∼8 Mb deletion on chromosome 8 (**Fig. S5D**), whereas in Ideo, 88.14% of 1,097 genes from 1,056 orthogroups corresponded to loss of the right arm of chromosome 14 (**Fig. S5G**).

At the gene-copy level, core genes constituted the largest fraction of each genome (∼55–86%), while soft-core and shell genes together accounted for most accessory gene copies (95–99%) (**Fig. 2B**; **Table S5**). Rare and private orthogroups contributed smaller proportions, although some variation was observed across cultivars. For example, YY5 v2.0 and CDC Bethune v3.0 contained 1,754 and 2,013 private genes, respectively.

Although total gene numbers differed among genomes, these differences were largely attributable to variation in accessory gene categories rather than the conserved core component. Fiber flax and linseed cultivars showed largely similar core gene complements, indicating that the fundamental gene repertoire of cultivated flax remains well-defined. Functional annotation patterns further supported this observation: approximately 55% of core genes were associated with GO annotations, whereas only ∼3.6% of private genes received GO annotations (**Table S5**), suggesting that the functional annotation of lineage-specific genes remains sparse.

Pangenome accumulation analysis further revealed that the total number of orthogroups increased with the addition of genomes, from ∼49,000 to >72,000, indicating continued discovery of novel gene orthogroups with increased genomic diversity, whereas the core genome decreased from ∼49,000 to ∼31,000 orthogroups, reflecting increasing gene presence–absence variation among cultivars (**Fig. 2C**).

### Phylogenetic analysis of the pangenome

A maximum likelihood phylogenomic tree was constructed using 20,178 single-copy orthologous genes across 17 cultivated flax genomes plus the wild progenitor genome *L. bienne* (LIN1917) (**Fig. 2D**). The wild accession was resolved as basal to cultivated flax, while cultivated *L. usitatissimum* accessions formed several well-supported clades. Fiber flax cultivars were not recovered as a monophyletic group but were distributed across multiple clades. Dual-purpose cultivar K-3018 was positioned between linseed accessions and the fiber flax accession Atlant, while Gaosi grouped with the fiber flax accessions.

### Morphotype-associated presence–absence variation

Presence–absence variation (PAV) analysis was conducted using nine linseed and six fiber flax cultivars. The two dual-purpose cultivars (K-3018 and Gaosi) were excluded from the statistical enrichment analysis to avoid ambiguity in morphotype assignment but were retained in downstream visualizations to evaluate their relationships with the two primary morphotypes. Using a tiered classification framework, 632 orthogroups met the predefined Tier 1 or Tier 2 enrichment criteria and displayed morphotype-biased presence–absence patterns (**Table S6**; **Fig. S6**), including 310 Tier 1 candidates showing the strongest contrasts between morphotypes and 322 Tier 2 candidates identified using more relaxed occurrence thresholds. Among these candidates, 432 orthogroups were enriched in linseed and 200 were enriched in fiber flax.

Heatmap visualization of all 17 cultivars, including the two dual-purpose accessions, revealed that K-3018 and Gaosi exhibited intermediate presence–absence patterns between the linseed and fiber flax groups (**Fig. S6**). However, K-3018 shared a larger proportion of the linseed-associated orthogroup profile, whereas Gaosi displayed a more mixed pattern.

Despite the large number of tiered candidates, none remained significant after Benjamini–Hochberg multiple-testing correction (FDR > 0.05). These results suggest that while numerous orthogroups exhibit modest morphotype-associated presence–absence differences, large-effect morphotype-specific gene content variation is limited among cultivated flax genomes. Collectively, the results indicate that gene presence–absence variation contributes relatively little to fiber flax–linseed divergence compared with the sequence-level variation identified through population genomic analyses.

### Repeat landscape of the flax pangenome

Repeat annotation using a flax pangenome TE library (You et al., 2026) revealed highly consistent repeat composition across genomes. The newly assembled genomes of Bison, Novelty, and Laura contained ∼65% repetitive sequences, comparable to CDC Bethune v3.0, with Class II DNA transposons representing the dominant component (∼36–37%), exceeding Class I retrotransposons (∼14–15%) (**Fig. 3A**; **Table S7**).

**Fig. 3.**
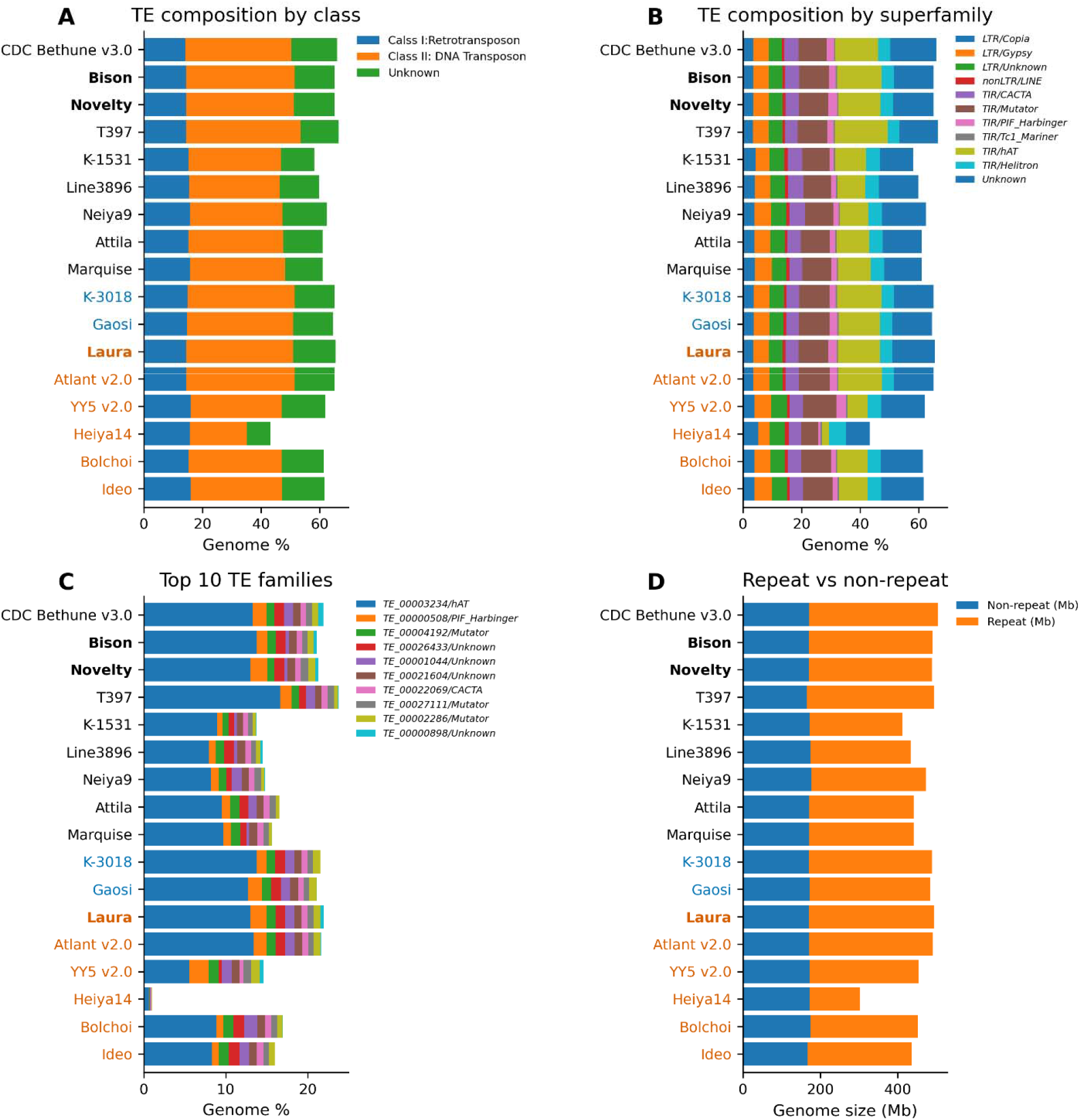
Comparative analysis of transposable elements across flax genomes. **A** Proportion of major TE classes across cultivars, showing dominance of Class II DNA transposons over Class I retrotransposons. **B:** TE composition by superfamily, highlighting the high abundance of *hAT* elements relative to LTR retrotransposons. **C:** Genome proportions of the ten most abundant TE families, with the *hAT* family *TE_00003234* as the predominant repeat across all cultivars. **D:** Comparison of repeat and non-repeat genome fractions. Cultivar names are shown in black for linseed, blue for dual and orange for fiber flax.

Across the pangenome, retrotransposon content remained relatively stable (∼14–16%), whereas variation in total repeat content was primarily associated with differences in DNA transposon abundance (**Fig. 3A**). Most cultivars exhibited high DNA transposon content (∼30–39%), while Heiya14 had fewer due to its smaller and incomplete assembly.

At the superfamily level, *hAT* elements were the most abundant DNA transposons across all cultivars, followed by *Mutator* elements (**Fig. 3B**). LTR retrotransposons (*Copia* and *Gypsy*) represented a smaller and relatively uniform fraction of the genome. Among individual TE families, the *TE_00003234 hAT* family alone accounted for ∼13–15% of the genome in most cultivars but was nearly undetectable in Heiya14 (**Fig. 3C**).

Comparison of repeat and non-repeat fractions showed that the latter was consistent across genomes at ∼170 Mb, whereas repeat content varied more across genomes (**Fig. 3D**). These results indicate that genome size variation among cultivars is primarily driven by differences in DNA transposon content, particularly *hAT* elements, although some of the observed variation may also reflect unresolved assembly challenges in highly repetitive regions.

### Genome-wide structural variation across flax genomes

Genome-wide structural variation (SV) was assessed using SyRI by comparing each assembly to the CDC Bethune v3.0 reference. Chromosome-scale organization was largely conserved for Bison, Novelty and Laura, with most aligned regions being classified as syntenic (**Fig. 4**). Small-scale SVs, including localized inversions and minor intrachromosomal rearrangements, were found to be enriched in low-gene-density regions.

**Fig. 4.**
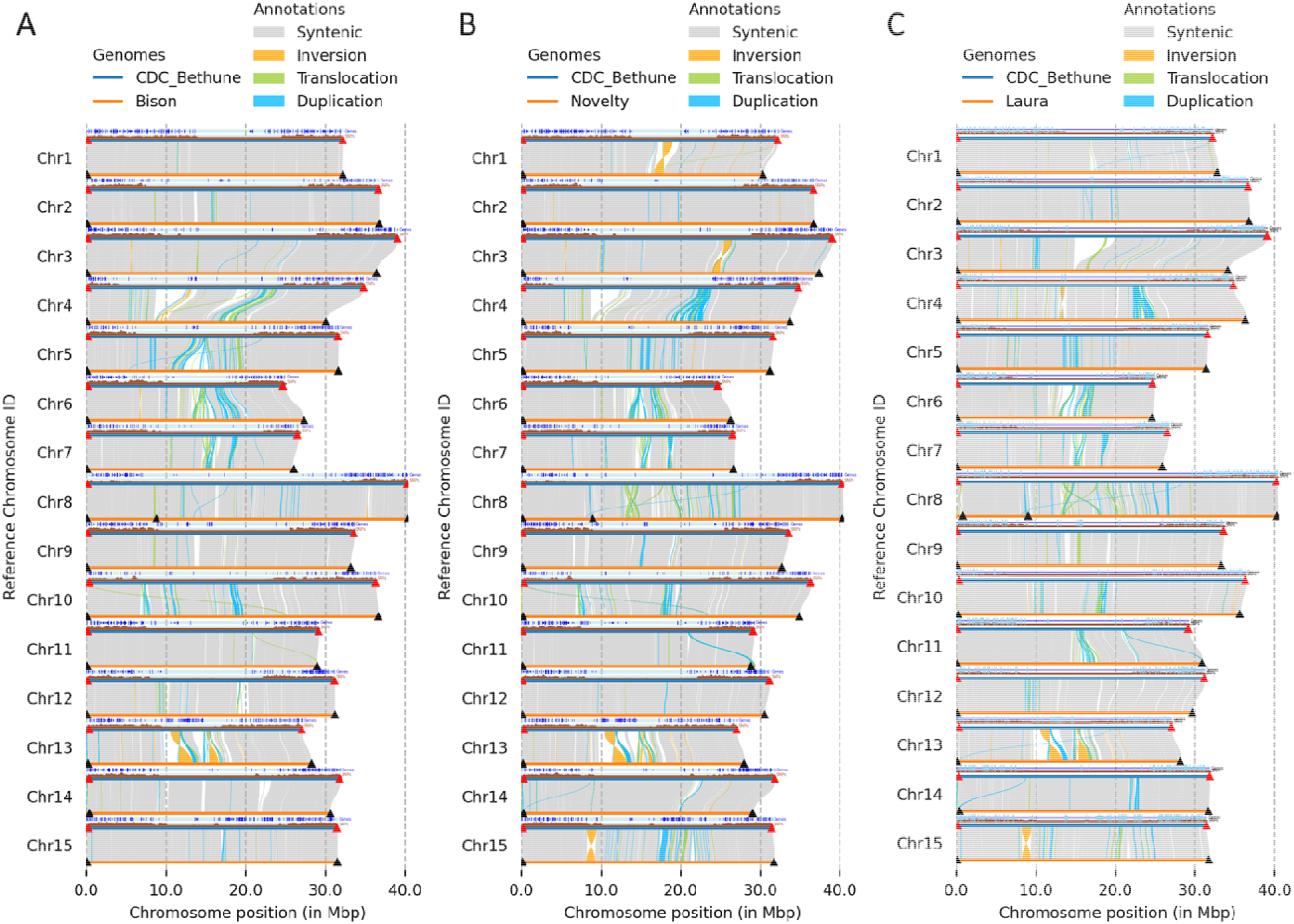
Genome-wide structural variation between CDC Bethune 3.0 and the three cultivars sequenced in this study revealed by SyRI-based comparative analysis. **A:** Bison. **B:** Novelty. **C:** Laura. Syntenic regions, inversions, translocations, and duplications are shown along chromosomes 1–15. Triangles indicate the positions of telomeric repeats detected at chromosome termini.

Comparisons with previously published assemblies showed generally high collinearity, particularly for Atlant v2.0 and Gaosi, which closely matched CDC Bethune v3.0 and the newly assembled genomes, with only minor differences such as small inversions on chromosomes 13 and 15 (**Figs. S5A,B**). In contrast, several assemblies exhibited large-scale structural variations (**Figs. S5C-H**). For example, YY5 v2.0 showed a ∼10 Mb inversion on chromosome 13 and additional inversions on chromosomes 2 and 10. Substantial variations in pseudomolecule structure were also observed in Line3896 and Heiya14, prompting reassembly of these genomes to assist structure comparison (**Figs. S5K-L**). K-3018 (**Fig. S5J**) and K-1531 (**Fig. S5I**), published as contigs, were also scaffolded to chromosomes using CDC Bethune v3.0 as a reference.

Some large rearrangements were not supported by telomeric repeat positions or were in low-gene-density or pericentromeric regions such as the inversion of the central region on chromosome 2 of YY5 v2.0 (**Fig. S5C**). Large terminal unaligned regions were also observed, including an ∼8 Mb region on the left arm of chromosome 8 in T397 and loss of the right arm of chromosome 14 in Ideo (**Figs. S5D** and **S5G**). Therefore, we treated these features as potential assembly or scaffolding artifacts and excluded them from SV analysis.

After exclusion of these regions, all remaining SVs ≥50 bp were retained for downstream analysis (**Fig. 5**). Across cultivars, particularly Bison, Novelty, Laura, Atlant v2.0, and Gaosi, SV profiles were extensively similar, with deletions representing the most frequent class, followed by duplications and translocations (**Fig. 5A**). However, duplications and translocations contributed a larger proportion of total SV span (**Fig. 5B**).

**Fig. 5.**
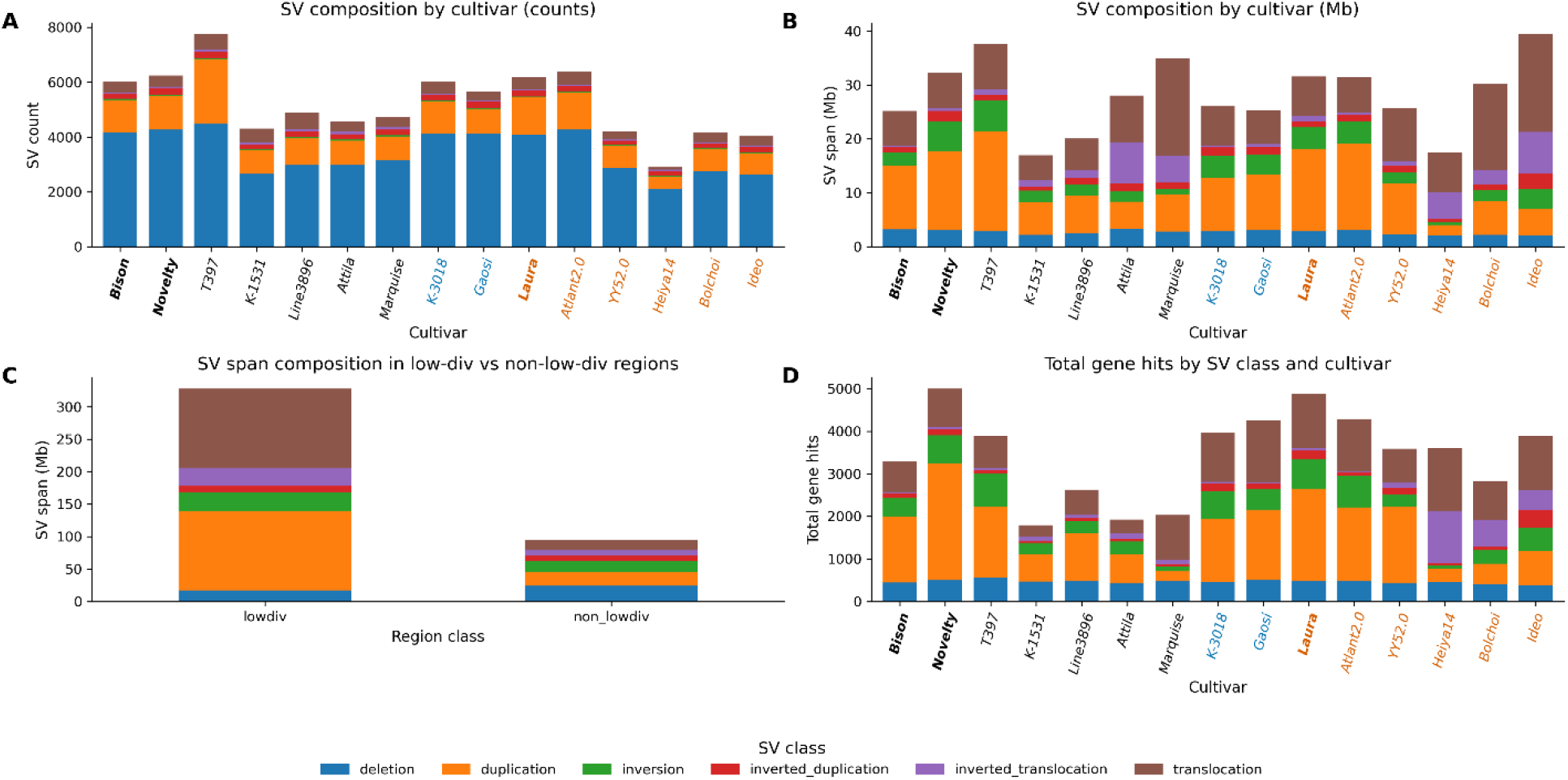
Genome-wide structural variation across the flax pangenome. **A:** Number of structural variants (SVs) per cultivar, partitioned by SV class. **B:** Total genomic span of SVs by cultivar and SV class. **C:** Distribution of SV span across low-diversity (pericentromeric) and non-low-diversity regions. **D:** Number of genes overlapping SVs by cultivar and SV class. Cultivar names are shown in black for linseed, blue for dual and orange for fiber flax.

SVs were enriched in low-diversity, repeat-rich regions (**Fig. 5C**), where duplications and translocations accounted for most of the affected sequence. In contrast, non-low-diversity regions showed fewer and smaller SVs. Analysis of gene overlap indicated that duplications and translocations accounted for most gene-associated SVs, whereas deletions affected fewer genes per event compared with duplications and translocations (**Fig. 5D**). These patterns indicate that structural variation is unevenly distributed across the genome, with larger events concentrated in repeat-rich regions.

### Genetic differentiation between fiber flax and linseed

Because no morphotype-specific genes were retained after multiple testing in PAV analysis, population genomic analyses were performed using 407 flax accessions from the flax core collection (94 fiber flax and 313 linseed accessions) to assess sequence-level differentiation.

Principal component analysis (PCA) based on ∼1.7 million genome-wide SNPs revealed clear population structure (**Fig. 6A**). The first two principal components (PC1 and PC2) largely separated the two morphotypes, with fiber flax accessions showing a more compact distribution while linseed accessions were more dispersed, with partial overlap between groups indicating shared genetic background.

**Fig. 6.**
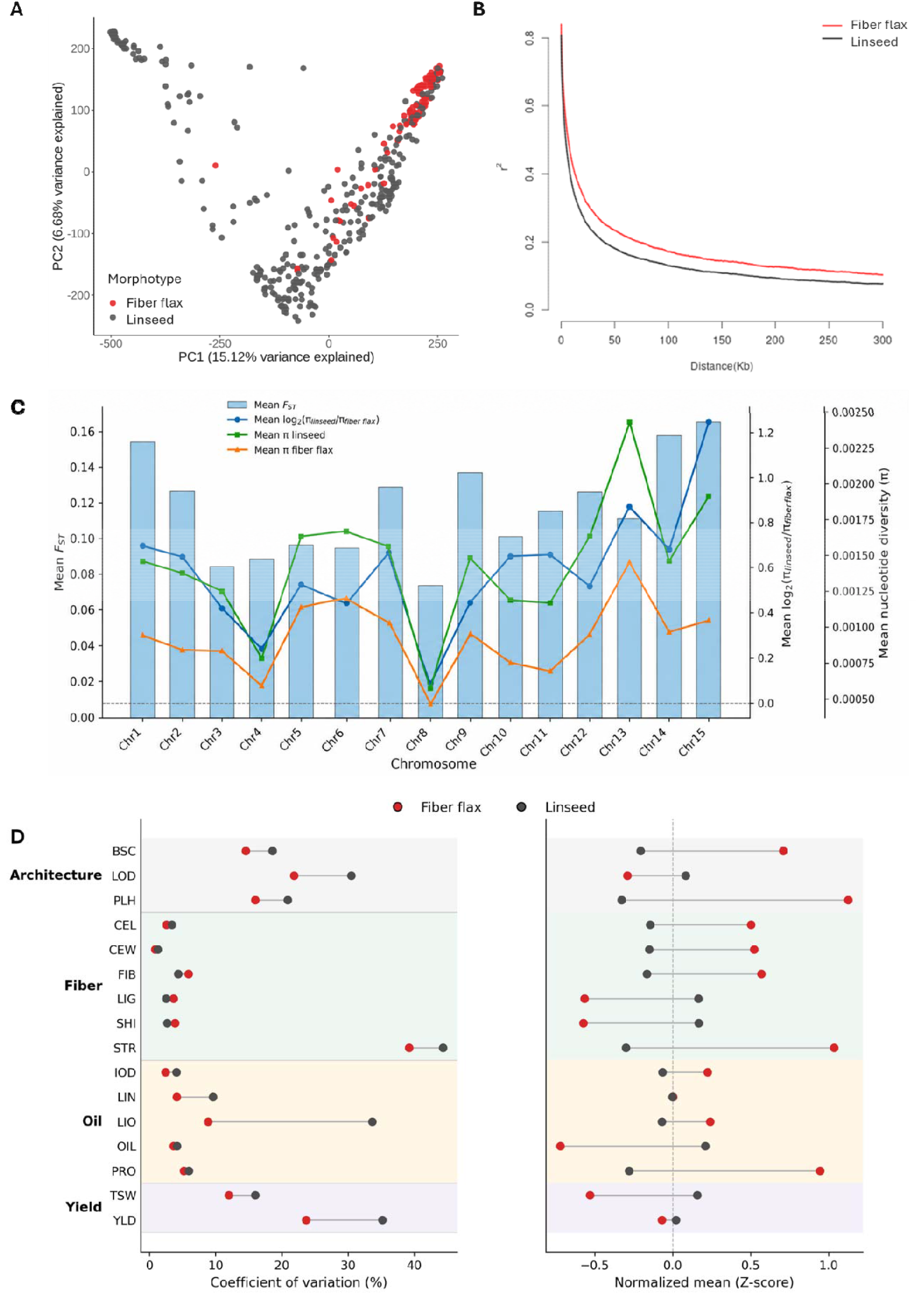
Population genomic differentiation and chromosome-scale diversity patterns in fiber flax and linseed. **A:** Principal component analysis (PCA) based on genome-wide SNP variation showing population structure between fiber flax and linseed accessions. **B:** Genome-wide linkage disequilibrium (LD) decay. LD (*r*^2^) was estimated from 313 linseed and 94 fiber flax accessions. LD was calculated from filtered SNPs (MAF ≥ 0.05, call rate ≥ 0.80, heterozygosity ≤ 0.20) and summarized up to 300 kb using PopLDdecay(Zhang et al., 2019). **C:** Chromosome-level summary of population differentiation and nucleotide diversity. Bars represent mean *F_ST_* per chromosome. Lines show mean log₂(*π*_linseed_/*π*_fiber fla*r*_) and mean nucleotide diversity (n) for linseed and fiber flax populations. **D:** Trait-level comparison of phenotypic variation and trait performance between morphotypes. Normalized mean values (Z-scores) between morphotypes illustrate directional shifts in trait performance associated with morphotype-specific selection. Trait abbreviations: BSC, branching score; LOD, lodging score; PLH, plant height (cm); CEL, cellulose content (%); CEW, cell wall content (%); FIB, fiber content (%); LIG, lignin content (%); SHI, shive content (%); STR, straw weight (g); IOD, iodine value; LIN, linolenic acid content (%); LIO, linoleic acid content (%); OIL, oil content (%); PRO, protein content (%); TSW, thousand-seed weight (g); YLD, seed yield (t/ha).

Linkage disequilibrium (LD) decay differed between morphotypes (**Fig. 6B**). LD decayed to *r*^2^ = 0.2 at 72.8 kb in fiber flax and 39.8 kb in linseed, and to *r*^2^ = 0.1 at ∼300 kb and 176.4 kb, respectively. Thus, LD persisted approximately 1.7–1.8 times longer in fiber flax, indicating more extended haplotype blocks.

Genome-wide nucleotide diversity (*π*) was lower in fiber flax. The window-weighted mean *π* was 8.99 x 10^—4^in fiber flax and 1.38 x 10^—3^in linseed, corresponding to a *π* ratio of ∼0.65 and representing a ∼35% reduction. Site-based estimates showed a similar pattern (fiber flax: 0.2289; linseed: 0.3444), and reduced diversity was observed across all chromosomes (**Figs. S7B-C**), indicating lower effective diversity in fiber flax.

The *π*_*fiber flax*_/*π*_*linseed*_ ratio showed that most windows clustered around 0.6–0.7, with 59.5% of windows <0.8 and 29.3% showing similar diversity between morphotypes (0.8-1.2) (**Fig. S7D**). The reciprocal ratio (*π*_*linseed*_/*π*_*fiber flax*_) exceeded one across most of the genome, with large window peaks showing markedly elevated values (**Fig. S7E**), indicating regions of stronger diversity reduction in fiber flax.

Chromosome-level summaries of nucleotide diversity revealed a consistent reduction of diversity in the fiber flax population across all chromosomes (**Fig. 6C; Table S8**). Mean *π* values ranged from 0.00078 to 0.00243 in linseed and from 0.00047 to 0.00145 in fiber flax, resulting in *π* ratios (*π*_*linseed*_/*π*_fiber_) exceeding one for all chromosomes (1.15–3.00). However, the magnitude of divergence varied among chromosomes, with the strongest reductions observed on Chr15, Chr1, and Chr14, whereas weaker divergence was observed on Chr8 and Chr4. The high coefficients of variation (*CV*) in *π*_*linseed*_/*π*_fiber_ ratios (*CV* = 43–337%) indicate considerable heterogeneity within chromosomes, suggesting that diversity reduction is driven by localized selective sweeps rather than uniform genome-wide processes. These patterns are consistent with chromosome-specific variation in *F_ST_* (**Fig. S7A**), reinforcing the conclusion that morphotype divergence is concentrated in discrete genomic regions.

### Phenotypic variation and trait divergence between fiber flax and linseed

To assess the impact of long-term selection on phenotypic variation, we compared trait means and coefficients of variation (CV) between fiber flax and linseed populations across 16 breeding-oriented agronomic and fiber traits (**Fig. 6D**).

Across all traits, linseed exhibited consistently higher or comparable phenotypic variation relative to fiber flax, with no evidence of systematic reduction in CV in the fiber flax population. This pattern closely mirrors genome-wide nucleotide diversity patterns, indicating that domestication and morphotype divergence have not resulted in a substantial loss of overall genetic variation.

Despite similar levels of phenotypic variation, pronounced differences in trait means were observed between morphotypes for multiple traits. Fiber-associated traits, including cellulose content (CEL), cell wall content (CEW), and straw yield (STR), showed elevated mean values in fiber flax, whereas oil-related traits such as oil content (OIL), iodine value (IOD), and thousand seed weight (TSW) were higher in linseed. Notably, several traits (e.g., CEL, CEW, shive content (SHI), IOD, OIL, protein content (PRO), and STR) exhibited comparable CV between morphotypes but markedly divergent means, suggesting that directional selection has primarily acted on trait means without markedly altering the underlying variance.

### Morphotype-associated quantitative trait nucleotides (QTNs)

To characterize the genetic architecture of morphotype-related traits without sample size bias, QTNs were identified in a combined diversity panel of 393 linseed and 85 fiber flax accessions for which phenotypic data were available. The analyses were conducted using five multi-locus models implemented in the mrMLM package (Zhang et al., 2020b) Genome-wide association study (GWAS) analyses were performed using BLUP values derived from multi-environment trials (four years × two locations). A total of 2,042 QTN signals were initially detected, which were consolidated into 1,712 unique trait–SNP associations after removing redundant detections across models. The number of QTNs varied among traits, ranging from 61 to 181 per trait (**Fig. 7A**, **Table S9**), with generally higher counts observed for fiber and oil-related traits.

**Fig. 7.**
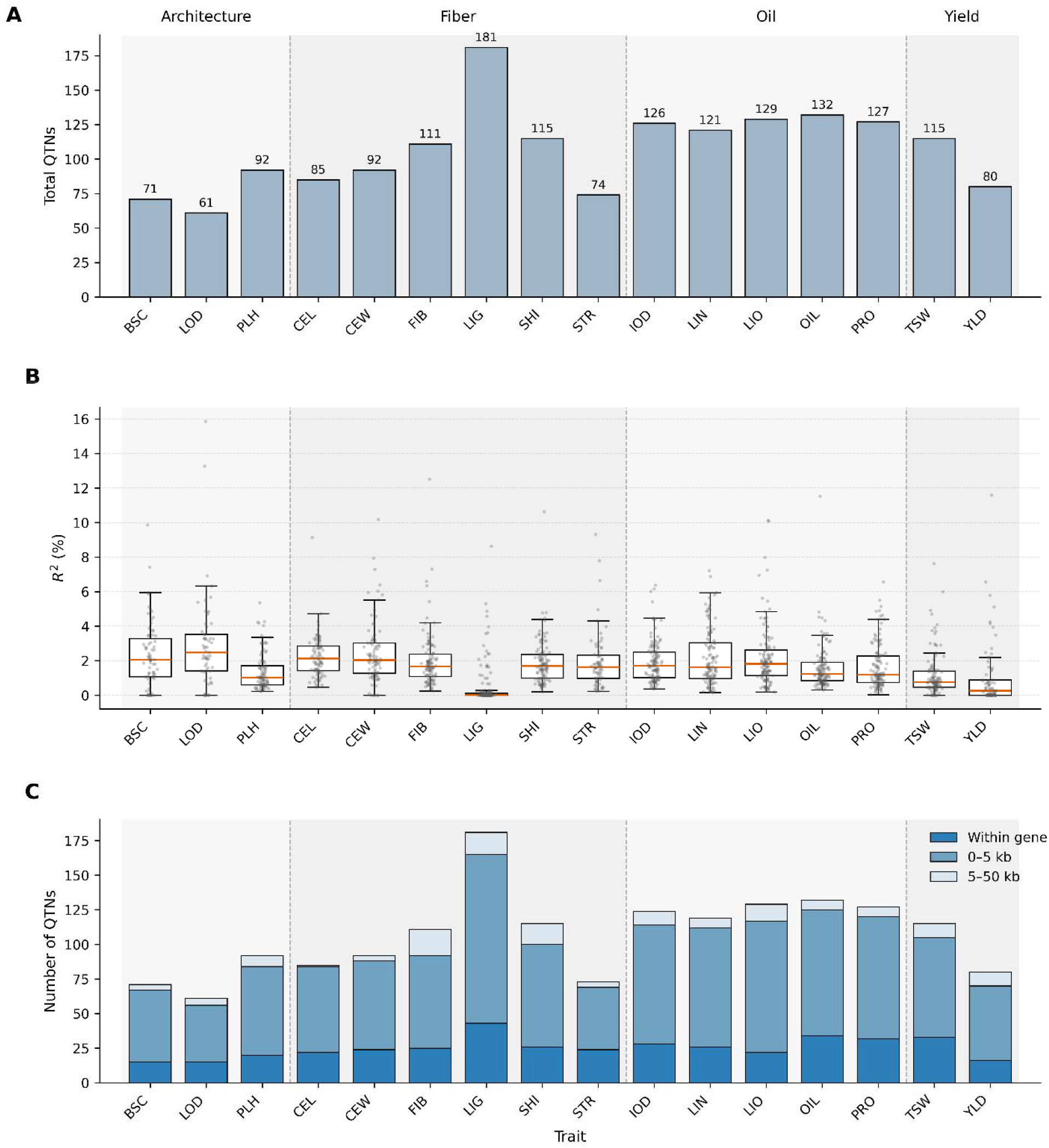
Genetic architecture of 16 morphotype-related traits inferred from quantitative trait nucleotide (QTN) analysis of the whole diversity panel. QTNs were identified using a combined panel comprising 393 linseed and 85 fiber flax accessions. **A:** Number of unique QTNs detected for each trait, grouped into architecture, fiber, oil, and yield categories. **B:** Distribution of QTN effect sizes for each trait, expressed as the proportion of phenotypic variance explained (*R*²). dots represent individual QTNs. **C:** Genomic distribution of QTNs relative to annotated genes. QTNs were classified into mutually exclusive categories based on distance to the nearest gene: within genes, and 0–5 kb, or 5–50 kb from nearest gene. Stacked bars show the number of QTNs in each category for each trait. Trait abbreviations: BSC, branching score; LOD, lodging score; PLH, plant height (cm); CEL, cellulose content (%); CEW, cell wall content (%); FIB, fiber content (%); LIG, lignin content (%); SHI, shive content (%); STR, straw weight (g); IOD, iodine value; LIN, linolenic acid content (%); LIO, linoleic acid content (%); OIL, oil content (%); PRO, protein content (%); TSW, thousand-seed weight (g); YLD, seed yield (t/ha).

Effect-size distributions indicated that most QTNs explained a small proportion of phenotypic variance, consistent with a polygenic architecture (**Fig. 7B**, **Table S9**). Median *R*² values were typically low (generally <2%), with only a small number of loci exhibiting moderate effects, indicating that trait variation is largely controlled by many small-effect loci.

Analysis of genomic distribution showed that QTNs were strongly enriched in gene-proximal regions (**Fig. 7C**, **Table S9**). Of the 1,712 unique QTNs, 405 (23.7%) were located within genes and 1,163 (68.0%) within 0–5 kb of annotated genes, whereas only 139 (8.1%) were located 5–50 kb away and just five QTNs (<0.3%) were found beyond 50 kb. These QTN patterns provide a basis for integrating association signals with population genomic analyses to identify candidate regions underlying morphotype divergence (**Fig. 8**).

**Figure 8.**
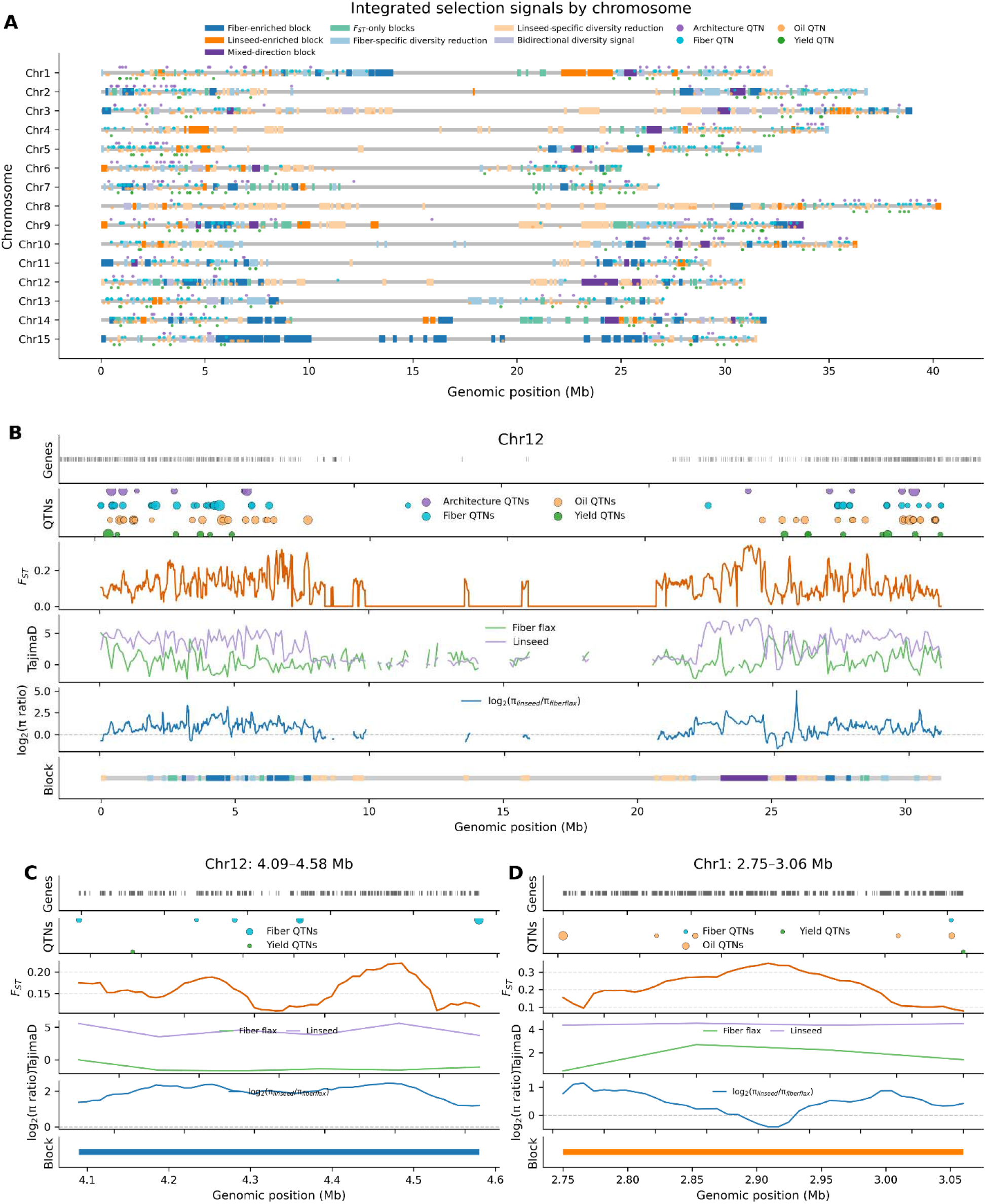
Integrated selection signals and candidate genomic regions underlying divergence between fiber flax and linseed flax morphotypes. **A**: Genome-wide distribution of integrated selection signals across chromosomes. Grey horizontal lines represent chromosomes. Colored segments indicate merged genomic blocks (superblocks) inferred from the integration of multiple metrics, including fiber flax-enriched (blue), linseed-enriched (orange), mixed-direction (purple), and *F_ST_*-only blocks (green), as well as blocks supported primarily by nucleotide diversity contrasts (*π*-only signals). Dots represent individual QTNs detected from the whole diversity panel, colored by trait categories. **B:** Chromosome-scale view of integrated selection signals along Chr12. Tracks (top to bottom) show gene annotations, QTN positions, windowed *F_ST_*, Tajima’s D for fiber flax and linseed populations, nucleotide diversity ratio log₂(*π*_*linseed*_/*π*_*fiber flax*_), and the inferred block classification. **C, D:** Representative regions illustrating integrated selection signals. **C:** fiber flax-enriched block (Chr12: 4.09-4.58 Mb), **D**: linseed-enriched block (Chr1:2.75–3.06 Mb). For each region, tracks are displayed in the same order as in (**B**), highlighting the concordance among differentiation, diversity, and QTN signals.

### Integrated identification of morphotype-enriched genomic blocks

To better resolve genomic regions associated with fiber flax–linseed divergence, we integrated population genetic signals, GWAS-derived QTNs, and gene annotations within a sliding-window framework. Genome-wide profiles of genetic differentiation (*F_ST_*), nucleotide diversity differences (log₂(*π*_*linseed*_/*π*_*fiber flax*_)), and Tajima’s D were combined with morphotype-specific QTNs to define candidate regions. Windows exceeding empirical thresholds (top 10% for *F_ST_* and diversity signals (*F_ST_* > 0.20; *π*_*linseed*_/*π*_*fiber flax*_> 2.92; *π*_*fiber flax*_/*π*_*linseed*_ > 1.10); SNP count ≥20) were merged into larger contiguous regions, hereafter referred to as morphotype-enriched genomic blocks (**Fig. 8A**; **Tables S10-S11**).

Across the genome, these integrated signals were unevenly distributed and clustered in discrete chromosomal regions (**Fig. 8A**). In total, 590 genomic blocks were identified across all signal classes. Among *F_ST_* -supported regions, 87 blocks were classified as fiber flax-enriched, 46 as linseed-enriched, 18 as mixed-direction, and 88 *F_ST_*-only blocks. In addition, *π*-only blocks included 128 fiber flax-enriched, 204 linseed-enriched, and 19 mixed-direction regions that were supported primarily by directional diversity signals but did not exceed the *F_ST_* threshold (**Tables S10-S11**).

Fiber flax-enriched blocks spanned 33.85 Mb of the genome and contained 2,331 genes, whereas linseed-enriched blocks represented a smaller number of *F_ST_*-supported regions. Directional diversity patterns differed between block classes: fiber flax-enriched blocks showed elevated log₂(*π*_*linseed*_/*π*_*fiber flax*_), consistent with reduced nucleotide diversity in fiber flax, whereas linseed-enriched blocks showed reduced log₂(*π*_*linseed*_/*π*_*fiber flax*_), consistent with reduced diversity in linseed. Mixed-direction blocks spanned 9.73 Mb and contained signals from both morphotypes, while *F_ST_*-only blocks spanned 16.03 Mb and showed elevated differentiation without strong directional diversity shifts.

*π*-only blocks represented a substantial proportion of candidate regions, particularly in linseed, where 204 blocks spanned 43.65 Mb. Although these regions did not exceed the genome-wide *F_ST_* threshold, many showed moderate differentiation, indicating that directional diversity shifts can identify additional candidate regions not captured by *F_ST_* alone.

Integration with GWAS results showed that multiple morphotype-enriched blocks overlapped with QTNs identified in fiber flax or linseed populations. Several blocks contained clustered QTNs and nearby candidate genes, suggesting that these regions may contribute to morphotype-associated trait variation. Although most QTNs had individually small effects (**Fig. 7B**), their local enrichment within candidate regions provides additional support for the biological relevance of these blocks.

Tajima’s D values differed between morphotypes within candidate regions (**Figs. 8B–D**). Genome-wide Tajima’s D values were generally higher in linseed than in fiber flax. Linseed-enriched regions frequently showed positive Tajima’s D values, whereas some fiber flax-enriched regions showed localized negative values. These contrasting patterns further distinguished candidate regions between morphotypes.

Representative regions illustrated the concordance among multiple signals (**Table S11**). A fiber flax-enriched region on Chr12 (4.09–4.58 Mb) showed elevated *F_ST_*, increased log₂(*π*_*linseed*_/*π*_*fiber flax*_), and reduced nucleotide diversity in fiber flax (**Fig. 8C**). Conversely, a linseed-enriched region on Chr1 (2.75–3.06 Mb) showed elevated *F_ST_* together with reduced log₂(*π*_*linseed*_/*π*_*fiber flax*_), consistent with reduced nucleotide diversity in linseed (**Fig. 8D**). In both examples, Tajima’s D differed between morphotypes, and QTNs or nearby genes were detected within the candidate intervals.

Overall, the integrated analysis identified discrete genomic blocks where population differentiation, directional diversity shifts, Tajima’s D, and QTN signals converge, revealing candidate regions for morphotype-associated divergence between fiber flax and linseed.

## DISCUSSION

### High-quality genome assemblies establish a reliable framework for flax comparative genomics

High-quality genome assemblies are essential for accurate pangenome and structural analyses, particularly in repeat-rich plant genomes (Nurk et al., 2022; Rhie et al., 2021). In this study, the three newly generated near T2T assemblies, together with CDC Bethune v3.0 (You et al., 2026), Atlant v2.0 (Dmitriev et al., 2020), and Gaosi (Lu et al., 2025), provide a consistent and reliable genomic framework for flax. These assemblies exhibit highly concordant chromosome structures, repeat composition, and gene completeness, supporting their use as a reference backbone for comparative analyses.

In contrast, several previously published assemblies display large-scale discrepancies, including megabase-scale inversions and terminal sequence loss that lack support from telomeric repeat organization or gene distribution. These inconsistencies are often localized in repeat-rich or low-gene-density regions and could reflect assembly or scaffolding limitations rather than true biological variation. Together, these observations highlight the importance of assembly quality and careful curation in pangenome analyses.

### Genome size variation is repeat-driven, whereas gene space remains conserved

Despite variation in genome size among flax cultivars (**Table 1**), gene content remains stable. The flax pangenome shares ∼170 Mb of non-repeat sequence (**Fig. 2D**), while high-confidence gene numbers and functional annotation proportions are highly consistent across cultivars. Core and soft-core gene families dominate the pangenome (**Table S6**), indicating that the functional gene repertoire is largely conserved across morphotypes and geographic origins.

In contrast, genome size variation is driven primarily by repeat expansion, particularly Class II DNA transposons. The predominance of *hAT* elements, especially the lineage-specific expansion of *TE_00003234*, distinguishes flax from many plant genomes in which LTR retrotransposons dominate genome expansion (Luo et al., 2017; Mascher et al., 2017; The International Brachypodium Initiative, 2010). Although TE dynamics contribute to genome architecture and pericentromeric organization (You et al., 2026), we found little evidence that TE variation directly underlies morphotype divergence.

### Morphotype divergence is primarily driven by allelic variation within a shared genomic background

A central question in flax domestication is whether fiber flax and linseed diverged through gene content differences or through allele-frequency shifts within a conserved genomic background. Pangenome analysis identified limited morphotype-associated presence–absence variation, and no orthogroups remained significantly associated with morphotype after multiple-testing correction. This pattern contrasts with several recently reported crop pangenomes in which morphotype or domestication-associated traits are accompanied by substantial gene presence–absence variation (Jayakodi et al., 2024; Jiao et al., 2025; Zhao et al., 2025), suggesting that flax diversification has been shaped primarily by allelic divergence within a conserved gene repertoire rather than by extensive gene-content variation.

In contrast, population genomic analyses revealed clear sequence-level differentiation between morphotypes. Fiber flax exhibited reduced nucleotide diversity, extended linkage disequilibrium, and a more compact population structure relative to linseed, consistent with stronger directional selection and a narrower genetic base (Hill and Robertson, 1966; Slatkin, 2008). Phylogenomic analysis further showed that fiber flax cultivars are not monophyletic, indicating that fiber-associated traits likely arose through repeated selection from a shared ancestral gene pool rather than from a single domestication lineage.

Genome-wide differentiation was highly heterogeneous, with most regions showing low to moderate divergence and a subset exhibiting elevated *F_ST_* and directional shifts in nucleotide diversity. Integration of *F_ST_*, *π* ratios, and QTN signals identified discrete morphotype-enriched genomic blocks distributed across chromosomes. Many candidate regions were supported primarily by directional diversity shifts, rather than extreme *F_ST_* values, consistent with adaptation through distributed allele-frequency changes rather than classic selective sweeps (Barrett and Schluter, 2008; Hermisson and Pennings, 2005; Jain and Stephan, 2018; Marques et al., 2018).

Patterns of Tajima’s D (Tajima, 1989) further support this interpretation. Linseed genotypes showed predominantly positive Tajima’s D values, whereas fiber flax genotypes displayed lower values with localized negative regions. These patterns are consistent with a model of selection on standing genetic variation observed in other crop species (Hufford et al., 2012; Meyer and Purugganan, 2013), in which multiple haplotypes contribute to adaptation and allele frequency shifts are moderate (Hermisson and Pennings, 2005; Messer and Petrov, 2013; Tajima, 1989).

The contrasting genetic architectures inferred from GWAS are also consistent with these morphotype-associated genomic patterns. Fiber-related traits were associated with fewer loci of moderate-to-large effect, whereas linseed traits showed a broader polygenic architecture involving many small-effect QTNs (Boyle et al., 2017). This difference is consistent with the stronger genetic bottleneck and reduced diversity observed in fiber flax populations. Notably, phenotypic variation remained comparable between morphotypes for many traits despite reduced nucleotide diversity in fiber flax, suggesting that directional selection altered trait means without substantially reducing overall phenotypic variability.

Together, these results support a model in which fiber flax represents a derived morphotype shaped primarily by directional selection acting on pre-existing allelic variation within a largely conserved genomic background.

### Integration of population genomics and association mapping identifies candidate regions underlying morphotype divergence

By integrating population genetic signals, nucleotide diversity patterns, Tajima’s D, and GWAS-derived QTNs (Nielsen, 2005; Vitti et al., 2013), we identified genomic regions where multiple independent lines of evidence converge. This framework transforms dispersed SNP-level signals into biologically interpretable candidate intervals. Morphotype-enriched genomic blocks were non-randomly distributed and frequently overlapped with morphotype-associated QTNs, supporting their biological relevance. Fiber flax- and linseed-enriched blocks displayed directional signatures consistent with population-specific selection, whereas mixed-direction blocks likely reflect more complex or overlapping selection histories. Representative regions showed strong concordance among differentiation, diversity shifts, Tajima’s D, and QTN enrichment, providing candidate targets for future functional analyses of flax domestication and breeding traits.

### Limitations and Implications for flax genomics and breeding

Several limitations should be considered. First, the number of high-quality assemblies per morphotype remains limited, reducing the power to detect subtle PAV. Second, structural variation analyses remain sensitive to assembly quality, particularly in repeat-rich regions. Third, GWAS-derived QTNs represent statistical associations that require functional validation. Despite these limitations, our integrated analyses consistently indicate that the genetic variation underlying fiber flax and linseed divergence resides largely within a shared genomic background. The predominance of allele-frequency divergence over gene content variation suggests that key alleles are broadly shared between morphotypes but differ in frequency. These findings highlight standing genetic variation as an important resource for flax improvement and suggest that breeding programs can effectively exploit existing allelic diversity through recombination and selection.

## Conclusion

This study presents a morphotype-resolved flax pangenome integrating high-quality genome assemblies and population-scale analyses. Despite substantial genome size variation driven by transposable elements, gene content is largely conserved and shows limited morphotype-specific differences. Instead, population genomic analyses reveal reduced diversity, extended linkage disequilibrium, and localized genomic differentiation in fiber flax. Integration of population genomic signals and association mapping identifies genomic regions where multiple lines of evidence converge. Importantly, the predominance of moderate-effect signals, *π* -driven differentiation, and non-extreme Tajima’s D patterns collectively support a model in which adaptation primarily involved soft selective sweeps acting on standing genetic variation. Together, these results support that the view that flax morphotype divergence is primarily driven by selection on standing genetic variation within a shared genomic background, providing a comprehensive framework for understanding flax evolution and for improving both fiber flax and linseed traits.

## MATERIALS AND METHODS

### Pangenome materials

In this study, three flax (*Linum usitatissimum*) cultivars were newly sequenced: Novelty (CN 97392 or PI 522557), Bison (CN 33399 or PGR 5050), and Laura (CN 18983). Seeds of these cultivars were obtained from Plant Gene Resources of Canada (PGRC, Saskatoon, Canada) and grown in a greenhouse. Seeds harvested from a single plant of each cultivar were used for genome sequencing to minimize heterogeneity. Bison is a linseed cultivar developed in 1930 by the North Dakota Experiment Station in the United States and registered in Canada in 1930, with resistance to Fusarium wilt and moderate resistance to powdery mildew. It is now used primarily as a foundational germplasm in disease-resistance studies. Novelty is a Canadian linseed cultivar registered in 1910, exhibiting moderate resistance to powdery mildew and susceptibility to Fusarium wilt (You et al., 2016). Laura is a Dutch fiber flax cultivar (Črepinšek and Kajfež-Bogataj, 2007).

In addition, 14 previously sequenced cultivars with near T2T or relatively complete genome assemblies were included in this study. In total, 17 assemblies representing the two major morphotypes, linseed and fiber flax, as well as dual-purpose type, were analyzed. The linseed group comprised nine cultivars: Bison, Novelty, CDC Bethune v3.0 (You et al., 2026), T397 (Yadav et al., 2025), Line3896 (Dvorianinova et al., 2023b), Neiya9 (Zhao et al., 2023), K-1531 (Dvorianinova et al., 2023a), Attila and Marquise (Demenou et al., 2025). The fiber flax group included six cultivars: Laura, YY5 (Sa et al., 2021), Atlant (Dmitriev et al., 2020), Heiya14, Bolchoi and Idéo (Demenou et al., 2025). In addition, the dual-purpose type cultivars K-3018 (Arkhipov et al., 2024) and Gaosi (Lu et al., 2025) were included. Among these cultivars, K-1531, Atlant, and Lin3896 are Russian cultivars; K-1531 belongs to a *crepitans*type (*Linum usitatissimum* convar. с*repitans* (Boenn.) Dumort), characterized by capsule dehiscence (Dvorianinova et al., 2023a). K-3018 originates from former Yugoslavia and represents an *intermedia* type (dual-purpose type), but it is cultivated as an oilseed. Attila, Marquise, Idéo and Bolchoi are French cultivars. YY5, Neiya9 and Heiya14 are Chinese cultivars, while T397 is an Indian linseed cultivar. Although K-3018 and Gaosi were classified as dual-purpose cultivars for visualization and comparative purposes, they were grouped with linseed flax in most downstream analyses because these cultivars are primarily used as oilseed flax and showed close genomic affinity to linseed accessions.

### DNA preparation for sequencing

High–molecular-weight (HMW) genomic DNA was isolated from young leaf tissue of greenhouse-grown seedlings. Plant growth conditions, dark treatment prior to sampling, tissue collection, and DNA extraction procedures followed the protocol described previously in You et al. (2026). Briefly, leaf tissue from 2–3 leaf stage seedlings was harvested, flash-frozen in liquid nitrogen, and processed using the NucleoBond HMW DNA Kit (Macherey-Nagel, Düren, Germany) according to the manufacturer’s instructions. DNA quality and integrity were assessed using PicoGreen quantification (ThermoFisher Scientific, Waltham, MA, USA) and pulsed-field gel electrophoresis. High-quality HMW DNA (15–30 μg per sample) was submitted to the Centre d’expertise et de services Génome Québec (Montréal, QC, Canada) for library construction and sequencing.

### Library construction and genome sequencing

SMRTbell libraries were constructed from HMW genomic DNA following the Pacific Biosciences SMRTbell® Prep Kit 3.0 protocol, as described in You et al. (2026). In brief, genomic DNA was sheared using a Megaruptor 3 system (Diagenode), followed by DNA damage repair, end repair, and SMRTbell adapter ligation. Libraries were size-selected using an AMPure PB bead–based protocol and sequenced on the PacBio Sequel II platform (SMRT Cell 8M) using Sequel II Sequencing Kit 2.0 chemistry. Sequencing runs employed 30-hour movie times with a 2-hour pre-extension and adaptive loading at a concentration of 80 pM.

### Genome assembly

PacBio HiFi reads were assembled using hifiasm v0.19.9 (Cheng et al., 2024) with default parameters, following the assembly strategy described in You et al. (2026). To eliminate organellar contamination, assembled contigs were aligned against the flax chloroplast reference genome (NCBI accession NC_002762) using BLASTN, and chloroplast-derived sequences were removed. Redundant haplotigs and high-coverage artifacts were filtered using purge_dups v1.2.6 (Guan et al., 2020). For assemblies incorporating long-read scaffolding, contigs were further scaffolded using LRscaf (Qin et al., 2019) with raw PacBio HiFi circular consensus sequencing (CCS) reads.

Chromosome-scale pseudomolecules were constructed using RagTag v2.1 (Alonge et al., 2022), with CDC Bethune 3.0 (You et al., 2026) as the reference genome. Assembly completeness was assessed using Benchmarking Universal Single-Copy Orthologs (BUSCO) (Simão et al., 2015) in genome mode with the embryophyta_odb10 dataset. Here, we defined “near telomere-to-telomere (near T2T) assemblies” as those where chromosomes may still contain one to five gaps, and telomeric repeats are identified for most chromosomes.

### Telomere identification

Telomeric regions were identified using the conserved plant telomeric repeat motif [TTTAGGG]n (Shakirov et al., 2022). Genomic regions containing at least 100 consecutive repeat units were defined as telomeric sequences, and the outermost coordinates of these repeat arrays were designated as chromosome telomere positions.

### Genetic maps

High-density genetic linkage maps were used to support chromosome-scale assembly validation and to investigate structural variation. The 702 biparental recombinant inbred line (RIL) population Bison × Novelty (BN) was analyzed as previously described (Cloutier et al., 2024). Both parental lines were sequenced and assembled in the present study, providing an opportunity to directly integrate high-quality parental genome assemblies with genetic map information for structural validation and comparative analysis.

Population development, plant growth conditions, DNA extraction, library preparation, and sequencing procedures followed previously established protocols (Cloutier et al., 2024). Illumina short reads from each individual were aligned to the CDC Bethune v3.0 reference genome (You et al., 2026) using BWA v0.6.1 (Jo and Koh, 2015), and SNP discovery was conducted using SAMtools v1.12 (Li and Durbin, 2009) within the AGSNP pipeline and its updated versions (Kumar et al., 2012; You et al., 2012; You et al., 2011).

High-quality SNPs were retained using the following filtering criteria: minor allele frequency > 0.05, heterozygosity call rate < 0.2, and genotype call rate > 80%. The genetic map was constructed using QTL IciMapping (Meng et al., 2015) following established procedures, including segregation distortion testing, redundancy reduction through marker binning, linkage group formation based on recombination frequency thresholds, and marker ordering using heuristic optimization algorithms. When necessary, linkage groups containing markers from multiple chromosomes were separated into distinct groups to ensure concordance with chromosome-scale assemblies.

### Genome size estimation

Genome sizes of the three newly sequenced cultivars were estimated using flow cytometry, following the procedure previously described (You et al., 2026).

### Repeat sequence annotation

A flax pangenome consensus transposable element (TE) library was previously constructed (You et al., 2026), integrating TE annotations from ten genotypes spanning six *Linum* species. This dataset included *L. bienne* accession LIN1917, *L. decumbens* accession LIN1754, *L. grandiflorum* accession LIN1530 (You et al., 2026), *L. tenue* (Gutierrez-Valencia et al., 2022), and *L. lewisii* (Innes et al., 2023) as well as *L. usitatissimum* cultivars CDC Bethune (You et al., 2026), Laura, Bison, Novelty, and Line3896 (Dvorianinova et al., 2023b). For each assembly, TEs were *de novo* identified using the Extensive De novo Transposable Element Annotator (EDTA) v2.0.0 pipeline (Ou et al., 2019) to create species- or genotype-specific TE libraries that were subsequently merged using the EDTA utility script make_panTElib.pl to generate a unified flax pangenome TE consensus library.

In this study, repeat annotation of the three newly assembled genomes, together with 14 previously published flax assemblies (17 cultivars in total), was performed using RepeatMasker v4.1.2 (Tarailo-Graovac and Chen, 2009) with the consensus TE library described above. The use of a single reference TE library and standardized annotation pipeline ensured consistent repeat classification and comparable estimates of TE content across all assemblies.

### Structural and functional gene annotation

The three newly assembled genomes and the 14 previously published flax cultivar assemblies were all processed using an identical and standardized pipeline. All genome assemblies were first repeat-masked using RepeatMasker v4.1.2 (Tarailo-Graovac and Chen, 2009). *Ab initio* gene prediction was conducted using AUGUSTUS v3.4.0 (Stanke et al., 2006) implemented within Braker v3.0.3 (Bruna et al., 2021; Hoff et al., 2016). Gene models were supported by extrinsic evidence derived from spliced RNA-seq alignments and a curated plant protein reference dataset. The protein reference dataset combined Viridiplantae protein sequences from OrthoDB (odb10) (Kriventseva et al., 2019) with previously published annotations from *Linum usitatissimum* and other *Linum*. Predicted gene models were processed using a standardized and curated workflow.

First, models lacking complete coding sequences (missing start or stop codons) or containing short coding regions with less than 17 amino acids were removed. Second, repeat-related genes were filtered by searching predicted proteins against the TREP database using DIAMOND (e-value ≤ 1e−05). Third, predicted proteins were queried against UniProtKB using DIAMOND (e-value ≤ 1e−05), and models without significant matches were excluded.

To classify gene confidence levels, predicted proteins were aligned to the NCBI non-redundant (nr) database. Gene models with alignment coverage ≥80% for both query and subject and e-value ≤ 1e−10 were designated as high-confidence (HC), and the remaining were classified as low-confidence (LC). Functional annotations were assigned based on the best-scoring database hit.

Functional annotation of predicted protein-coding genes was performed using eggNOG-mapper (Cantalapiedra et al., 2021) based on the eggNOG v5.0 database (Huerta-Cepas et al., 2019). Protein sequences from each assembly were queried against the precomputed orthologous groups to infer functional categories through orthology assignment. Gene Ontology (GO) terms and Kyoto Encyclopedia of Genes and Genomes (KEGG) pathway annotations were retrieved directly from eggNOG-mapper outputs. For each genome, annotation summaries including the number of genes with GO and KEGG assignments were compiled for comparative analyses across the flax pangenome.

### Orthogroup analysis and phylogenetic analysis

Orthogroup inference was conducted using OrthoFinder v2.5.5 (Emms and Kelly, 2019) based on protein-coding gene sets from 17 flax cultivars together with the *L. bienne* accession LIN1917 (You et al., 2026), which was used as an outgroup. Given the extensive gene duplication observed across flax genomes, loci for phylogenetic reconstruction were identified using a relaxed “near-single-copy” criterion where orthogroups were required to contain genes from all 16 cultivated genomes, and where at least 15 of the 17 genomes had to exhibit single-copy representation within the orthogroup. Orthogroups with widespread duplication beyond this threshold were excluded.

For orthogroups meeting these criteria, a single representative gene was selected per genome. In cases where only 1-2 genomes had paralogous gene copies, all possible single-copy combinations were evaluated. Candidate combinations were assessed using the corresponding OrthoFinder gene tree by calculating the sum of pairwise patristic distances among selected tips. The combination minimizing the total phylogenetic distance—representing the most consistent orthologous clustering—was retained. When ties occurred, the set with the greatest cumulative protein sequence length was selected to reduce potential biases from truncated gene models.

Representative protein sequences from retained near-single-copy orthogroups were aligned using MAFFT (Katoh and Standley, 2013), and individual alignments were concatenated to generate a supermatrix. Maximum-likelihood phylogenetic inference was performed using RAxML-NG v1.2.2 (Kozlov et al., 2019) under the LG+G+F substitution model with 1,000 bootstrap replicates.

### Genome structural variation analysis

Genome-wide structural variations (SVs) were identified through whole-genome pairwise comparisons using CDC Bethune v3.0 as the reference assembly. Prior to alignment, chromosome identifiers were standardized across all assemblies to ensure consistent chromosome correspondence. Our assemblies follow the chromosome numbering system established by the first complete flax consensus genetic map (Cloutier et al., 2012) and adopted in CDC Bethune v2.0 (You et al., 2018). However, chromosome numbering in recently published flax genomes from other research groups differed due to inconsistent naming conventions.

To harmonize chromosome identities and prevent artificial detection of inter-chromosomal rearrangements, a systematic renaming scheme was applied prior to alignment. The following conversions were used to map chromosome identifiers from previously published flax genomes to the consensus numbering system: Chr2 → Chr10, Chr3 → Chr11, Chr4 → Chr12, Chr5 → Chr13, Chr6 → Chr14, Chr7 → Chr15, Chr8 → Chr2, Chr9 → Chr3, Chr10 → Chr4, Chr11 → Chr5, Chr12 → Chr6, Chr13 → Chr7, Chr14 → Chr8, and Chr15 → Chr9. In each conversion, the first chromosome identifier corresponds to the numbering used in previously published flax genomes, whereas the second corresponds to the consensus chromosome numbering system adopted in this study.

Whole-genome alignments were generated using minimap2 2.30 (Li, 2021) with parameters optimized for assembly-to-assembly comparison. The resulting alignments were processed using SyRI 1.7.1 (Goel et al., 2019) to detect structural variants, including inversions, translocations, duplications, and large insertions/deletions. Structural rearrangements were visualized at the chromosome scale using plotsr 1.1.0 (Goel and Schneeberger, 2022).

### Pangenome analysis

Orthogroups inferred across the 17 cultivated flax genomes were classified into standard pangenome categories according to their prevalence among genomes (Golicz et al., 2016b). Orthogroups present in all genomes were defined as core, whereas those missing in only one genome were considered soft-core. Orthogroups detected in three to 15 of the genomes were assigned to the shell category, while those restricted to two genomes were classified as rare and those found in a single genome were defined as private. Gene copy numbers were summarized for each category and genome to quantify variation in conserved versus accessory gene content across the flax pangenome.

To identify orthogroups exhibiting consistent presence–absence differences between flax morphotypes, enrichment analysis was performed using only linseed (n = 9) and fiber flax (n = 6) cultivars. The two dual-purpose cultivars (K-3018 and Gaosi) were excluded from the enrichment analysis but were retained for visualization in downstream heatmaps. For each orthogroup, presence–absence frequencies were compared between morphotypes using Fisher’s exact test, and false discovery rates (FDR) were estimated using the Benjamini–Hochberg procedure (Benjamini and Hochberg, 1995). Orthogroups were further classified into two confidence tiers based on the consistency of morphotype-enriched occurrence patterns. Tier 1 orthogroups were defined as those present in at least seven of nine linseed cultivars and in no more than one of six fiber flax cultivars, or present in at least five of six fiber flax cultivars and in no more than one of nine linseed cultivars. Tier 2 orthogroups met the same presence thresholds in the enriched morphotype but were allowed to occur in up to two cultivars of the contrasting morphotype. Orthogroups assigned to Tier 1 were excluded from Tier 2. This hierarchical classification was used to identify candidate orthogroups exhibiting strong and consistent morphotype-associated presence–absence variation. Pangenome composition was visualized using bar plots summarizing orthogroup and gene counts per category, along with stacked bar plots illustrating gene composition for each genome. Pangenome accumulation curves were generated by randomly permuting the order of genome addition to evaluate pan-genome expansion and core-genome contraction dynamics.

### Flax diversity panel and population structure

A core diversity panel comprising 407 flax accessions from 38 countries, including 313 linseed and 94 fiber flax accessions (You et al., 2017) was used for genomic analysis. These accessions were resequenced using whole-genome short-read sequencing. SNPs were identified using a customized variant-calling pipeline (You et al., 2012), with CDC Bethune 3.0 (You et al., 2026) as reference. After filtering for minor allele frequency ≥ 0.05, heterozygosity ≤ 0.2, and call rate ≥ 0.8, approximately 1.7 million high-quality SNPs were retained. Detailed procedures are described in You et al. (2026).

Population structure was assessed using principal component analysis (PCA) based on genome-wide SNPs. PCA was performed using the PrincipalComponentsPlugin implemented in TASSEL 5.0 (Bradbury et al., 2007), and morphotype groups were visualized in the resulting reduced dimensional space.

### LD decay analysis

Linkage disequilibrium (LD) decay was estimated separately for fiber flax (n = 85) and linseed (n = 293) accessions using PopLDdecay 3.43 (Zhang et al., 2019). Genome-wide SNPs were filtered to retain high-quality variants with minor allele frequency (MAF) ≥ 0.05, genotype call rate ≥ 0.80, and heterozygosity ≤ 0.20. Pairwise LD (*r²*) was calculated and summarized in physical distance bins up to a maximum inter-marker distance of 300 kb. Short- and long-distance bin sizes were set to 10 bp (--bin1 10) and 100 bp (--bin2 100), respectively. Mean *r²* values were plotted against physical distance to visualize LD decay patterns.

### Genome-wide association study (GWAS)

Multi-locus genome-wide association analysis was conducted using the mrMLM 5.0.1 R package (Zhang et al., 2020b), which integrates multi-locus models including mrMLM (Wang et al., 2016), FASTmrEMMA (Wen et al., 2018), ISIS EM-BLASSO (Tamba et al., 2017), FASTmrMLM (Zhang and Tamba, 2018), and pLARmEB (Zhang et al., 2017). Out of 407 accessions in the flax core collection, 376 with complete phenotypic data were retained for GWAS, including 293 linseed and 85 fiber flax accessions. A total of 27 traits relevant to linseed and fiber flax production were evaluated for five years at two locations (You et al., 2017). Best linear unbiased prediction (BLUP) values were used for association analysis, including eleven linseed-related traits analyzed in the linseed panel and eight fiber-related traits in the fiber flax panel. Significant quantitative trait nucleotides (QTNs) were identified using a logarithm of odds (LOD) threshold of 3.0.

### Chromosome-scale differentiation

Genetic differentiation between fiber and linseed flax was quantified using Weir and Cockerham’s *F_ST_*, nucleotide diversity (*π*), and Tajima’s D, calculated with VCFtools (Danecek et al., 2011). Genome-wide patterns were estimated using sliding windows of 100 kb with a 10-kb step size for *F_ST_* and *π*, and non-overlapping 100-kb windows for Tajima’s D. Windows with low SNP density (< 20 SNPs in a 100-kb window) were excluded to minimize bias.

Genome-wide *F_ST_* and *π* ratio profiles were used to identify differentiated regions. Windows in the top 10% of the genome-wide distributions were selected and merged if overlapping or adjacent, generating a nonredundant set of candidate divergence blocks. To reduce artifacts associated with low polymorphism, *π* ratios were calculated only for windows containing at least 20 SNPs in both populations. Genome-wide distributions were visualized using Manhattan-style plots with cumulative chromosomal coordinates.

### Morphotype-enriched genomic block identification

Morphotype-enriched genomic regions were identified by integrating population genetic statistics and GWAS signals using a sliding-window framework. Genome-wide genetic differentiation (weighted *F_ST_*) and nucleotide diversity differences were estimated in fixed windows and summarized as log₂(*π*_*linseed*_/*π*_*fiber flax*_). Windows with insufficient variation (<20 SNPs) were excluded. Candidate selection signals were defined using empirical thresholds, where windows in the top 10% of the genome-wide distribution for *F_ST_* and log₂(*π*_*linseed*_/*π*_*fiber flax*_) were retained.

Directional signals were inferred based on the diversity ratio: windows with elevated log₂(*π*_*linseed*_/*π*_*fiber flax*_) indicate reduced diversity in the fiber flax population (fiber flax-associated), whereas windows with reduced values indicate reduced diversity in the linseed population (linseed-associated). Windows exceeding the *F_ST_* threshold but lacking strong directional signals were classified as *F_ST_* -only candidates. Adjacent or overlapping candidate windows were merged to define genomic regions (superblocks), and summary statistics (e.g., mean, and maximum *F_ST_* and log₂ ratios) were calculated for each block.

To further support candidate regions, morphotype-specific QTNs identified from multi-locus GWAS analyses were projected onto the genome and intersected with candidate windows and merged blocks. For each block, the number and average effect size (mean t^2^ of overlapping QTNs from fiber flax and linseed populations were calculated. Gene annotations were intersected with candidate blocks to quantify gene content. In addition, Tajima’s D was calculated separately for fiber flax and linseed populations and summarized within each block to provide complementary evidence of selection.

Blocks were classified into four categories based on the combination of supporting signals: fiber flax-enriched blocks (high *F_ST_* and elevated log₂(*π*_*linseed*_/*π*_*fiber flax*_)), linseed-enriched blocks (high *F_ST_* and elevated log₂(*π*_*fiber flax*_/*π*_*linseed*_)), mixed-direction blocks (both *π* ratios elevated), and *F_ST_* -only blocks (high *F_ST_* without directional *π* signals). In all cases, “high” and “elevated” refer to values exceeding the top 10% of the genome-wide distribution.

### Data availability

All raw sequencing data and genome assemblies generated in this study have been deposited in the National Center for Biotechnology Information (NCBI) database under the following accessions. The genome assembly of CDC Bethune is available under accession JAZBJT000000000 (BioProject: PRJNA1060737; BioSample: SAMN39244156). The genome assembly of Laura is available under accession JAYKZD000000000 (BioProject: PRJNA1061544; BioSample: SAMN39273365). The Bison assembly is available under accession JAYKZB000000000 (BioProject: PRJNA1061532; BioSample: SAMN39272935). The Novelty assembly is available under accession JAYKZC000000000 (BioProject: PRJNA1061538; BioSample: SAMN39272985). All associated raw sequence reads are accessible through the corresponding BioProject records. The genome sequences and annotations were also deposited in Zenodo at https://zenodo.org/records/20668184. Any additional information is available from the corresponding author upon request.

### Supplementary data

Supplementary data to this article can be found online at http://xxx.

## Acknowledgments

This work was conducted as part of the Total Utilization Flax GENomics (TUFGEN) project funded by Genome Canada and other stakeholders, the A-base project no. 1142 funded by Agriculture and Agri-Food Canada, and the Diverse Field Crop Cluster (DFCC) managed by Ag-West Bio Inc (J-003426). We acknowledge Tracey James and Sara Martin for assistance with flow cytometry analyses.

## Credit authorship contribution statement

**Frank M. You**: Conceptualization, Methodology, Resources, Funding acquisition, Original draft, Formal analysis; **Chunfang Zheng, Pingchuan Li**: Investigation; **Tara Edwards**: Investigation, Writing - Review & Editing; **Khalid Y Rashid, Scott D Duguid, Helen Booker**: Resources, Funding acquisition, Writing - Review & Editing; **Sylvie Cloutier**: Conceptualization, Methodology, Resources, Funding acquisition, Writing - Review & Editing.

