## Supplemental tables and figures for "Integrated pangenome and population genomics reveal selection on standing genetic variation driving fiber flax-linseed divergence"

**Table S1**. Read statistics and genome size estimation of three flax cultivars along with the reference genome CDC Bethune v3.0.

| **Cultivar** | **HiFi read size (Gb)** | **Total number of reads** | **Read coverage (X)** | **Average read length (bp)** | **Assembled size**  **(Mb)** | **Genome size estimates by K-mer analysis (K=17)** | **Genome size by flow cytometry (Mb)** |
| --- | --- | --- | --- | --- | --- | --- | --- |
| CDC Bethune v3.0^†^ | 28.57 | 2,350,375 | 52 | 12,155 | 504.00 | 529.01 | 581 |
| Bison | 21.58 | 1,879,312 | 41 | 11,483 | 490.06 | 510.74 | 568 |
| Novelty | 24.22 | 2,244,606 | 45 | 9,501 | 489.26 | 519.56 | 567 |
| Laura | 22.92 | 2,070,824 | 42 | 11,069 | 494.28 | 531.23 | 576 |

† Data from You et al. (2026).

**Table S2.** Chromosome sizes of three flax genome assemblies and the reference genome CDC Bethune v3.0 using PacBio HiFi reads.

| **Chromosome** | **CDC Bethune v3.0 (ref)** ^†^ | | **Bison** | | | | | **Novelty** | | | | | **Laura** | | | |
| --- | --- | --- | --- | --- | --- | --- | --- | --- | --- | --- | --- | --- | --- | --- | --- | --- |
|  | **No. of scaffolds** | **Size (bp)** | | **No. of scaffolds** | | **Size (bp)** | | | **No. of scaffolds** | | **Size (bp)** | | | **No. of scaffolds** | | **Size (bp)** |
| Chr1 | 2 | 32,352,746 | | 1 | 32,255,778 | | 3 | | | 30,483,443 | | 2 | | | 32,978,378 | |
| Chr2 | 1 | 36,797,904 | | 1 | 36,824,130 | | 1 | | | 36,877,162 | | 1 | | | 36,968,402 | |
| Chr3 | 1 | 39,172,353 | | 1 | 36,480,313 | | 1 | | | 37,570,511 | | 1 | | | 34,270,047 | |
| Chr4 | 1 | 34,927,835 | | 1 | 30,077,852 | | 3 | | | 33,886,325 | | 1 | | | 36,473,621 | |
| Chr5 | 2 | 31,717,593 | | 2 | 31,648,237 | | 1 | | | 31,347,906 | | 1 | | | 31,573,969 | |
| Chr6 | 1 | 25,011,845 | | 1 | 27,377,534 | | 1 | | | 26,452,595 | | 1 | | | 24,745,759 | |
| Chr7 | 3 | 26,806,467 | | 1 | 26,068,398 | | 3 | | | 26,751,805 | | 1 | | | 26,016,147 | |
| Chr8 | 1 | 40,448,020 | | 3 | 40,339,355 | | 2 | | | 40,495,006 | | 3 | | | 40,512,198 | |
| Chr9 | 3 | 33,717,904 | | 3 | 33,201,001 | | 2 | | | 32,872,215 | | 2 | | | 33,448,514 | |
| Chr10 | 4 | 36,426,562 | | 2 | 36,714,767 | | 3 | | | 35,072,801 | | 4 | | | 35,782,596 | |
| Chr11 | 3 | 29,269,339 | | 2 | 29,221,277 | | 5 | | | 29,382,747 | | 2 | | | 31,124,826 | |
| Chr12 | 2 | 31,253,724 | | 1 | 31,240,139 | | 2 | | | 30,703,757 | | 1 | | | 29,826,650 | |
| Chr13 | 2 | 27,152,020 | | 3 | 28,276,286 | | 3 | | | 28,105,563 | | 2 | | | 28,297,954 | |
| Chr14 | 5 | 31,933,221 | | 4 | 30,663,210 | | 7 | | | 29,183,855 | | 3 | | | 31,823,488 | |
| Chr15 | 2 | 31,567,185 | | 1 | 31,547,584 | | 2 | | | 31,893,932 | | 3 | | | 31,914,170 | |
| Chr total | 33 | 488,554,718 | | 27 | 481,935,861 | | 39 | | | 481,079,623 | | 28 | | | 485,756,719[S | |
| Un | 134 | 15,444,289 | | 76 | 8,119,563 | | 74 | | | 8,181,172 | | 119 | | | 8,527,348 | |
| Overall | 167 | 503,999,007 | | 103 | 490,055,424 | | 113 | | | 489,260,795 | | 147 | | | 494,284,067 | |

† Data from You et al. (2026); Un: unanchored scaffolds.

**Table S3.** Genetic map of the Bison x Novelty recombinant inbreeding line (RIL) population of 702 individuals

| **Chromosome** | **No of markers** | **Distance (cM)** | **Misaligned markers*** |
| --- | --- | --- | --- |
| Chr1 | 214 | 258.25 | 0 |
| Chr2 | 110 | 180.81 | 1 |
| Chr3 | 75 | 165.32 | 0 |
| Chr4 | 102 | 174.67 | 0 |
| Chr5 | 113 | 177.02 | 1 |
| Chr6 | 111 | 168.60 | 0 |
| Chr7 | 160 | 188.93 | 3 |
| Chr8 | 71 | 161.84 | 1 |
| Chr9 | 86 | 149.36 | 0 |
| Chr10 | 76 | 159.61 | 0 |
| Chr11 | 127 | 168.98 | 1 |
| Chr12 | 136 | 190.91 | 0 |
| Chr13 | 74 | 142.45 | 0 |
| Chr14 | 119 | 171.08 | 0 |
| Chr15 | 75 | 148.50 | 0 |
| Total | 1649 | 2,606.33 | 7 |

* Misaligned markers were assigned to one chromosome during SNP discovery but aligned to a different chromosome in the genetic map.

**Table S4.** Assembly and chromosome sizes across 17 flax assemblies.

| **Chr** | **CDC Bethune v3.0** | **Bison** | **Novelty** | **T397** | **K-1531^†^** | **Line3896^†^** | **Neiya9** | **Attila** | **Marquise** | K-3018† | **Gaosi** | **Laura** | **Atlant v2.0** | **YY5 v2.0** | **Heiya14^†^** | **Bolchoi** | **Ideo** | **Min** | **Max** | **Median** | **Mean** |
| --- | --- | --- | --- | --- | --- | --- | --- | --- | --- | --- | --- | --- | --- | --- | --- | --- | --- | --- | --- | --- | --- |
| 1 | 32.35 | 32.26 | 30.48 | 31.29 | 28.66 | 29.96 | 31.58 | 32.82 | 32.77 | 32.78 | 33.03 | 32.98 | 32.99 | 31.79 | 21.37 | 34.51 | 32.14 | 16.53 | 37.77 | 26.46 | 27.21 |
| 2 | 36.80 | 36.82 | 36.88 | 36.42 | 26.75 | 30.20 | 32.08 | 31.31 | 31.67 | 36.67 | 36.54 | 36.97 | 36.84 | 29.85 | 22.60 | 32.40 | 31.43 | 19.82 | 38.89 | 27.92 | 28.16 |
| 3 | 39.17 | 36.48 | 37.57 | 38.57 | 24.25 | 28.95 | 42.83 | 31.72 | 33.79 | 38.76 | 34.24 | 34.27 | 35.82 | 26.36 | 23.57 | 31.77 | 32.47 | 19.91 | 42.83 | 31.58 | 30.87 |
| 4 | 34.93 | 30.08 | 33.89 | 36.10 | 23.47 | 24.98 | 19.91 | 30.76 | 26.97 | 33.91 | 33.58 | 36.47 | 36.32 | 24.86 | 16.87 | 30.99 | 32.12 | 17.72 | 39.91 | 29.85 | 28.21 |
| 5 | 31.72 | 31.65 | 31.35 | 32.87 | 20.95 | 24.27 | 30.01 | 26.45 | 25.41 | 30.48 | 31.68 | 31.57 | 32.26 | 24.52 | 19.76 | 25.73 | 26.46 | 27.75 | 40.76 | 32.13 | 32.74 |
| 6 | 25.01 | 27.38 | 26.45 | 29.39 | 23.53 | 20.54 | 36.22 | 19.77 | 18.93 | 26.83 | 24.81 | 24.75 | 28.58 | 28.49 | 17.54 | 19.82 | 19.37 | 24.75 | 40.51 | 31.91 | 32.38 |
| 7 | 26.81 | 26.07 | 26.75 | 28.14 | 22.24 | 21.70 | 26.49 | 21.86 | 25.92 | 26.07 | 27.43 | 26.02 | 27.75 | 17.72 | 17.61 | 24.30 | 25.45 | 24.81 | 40.71 | 31.95 | 32.17 |
| 8 | 40.45 | 40.34 | 40.50 | 31.56 | 30.55 | 31.58 | 35.76 | 40.70 | 42.94 | 41.04 | 40.71 | 40.51 | 40.76 | 31.05 | 21.50 | 38.89 | 37.77 | 26.07 | 41.04 | 32.03 | 32.48 |
| 9 | 33.72 | 33.20 | 32.87 | 33.68 | 27.85 | 28.32 | 34.74 | 30.94 | 30.20 | 32.08 | 33.42 | 33.45 | 32.13 | 32.04 | 21.66 | 30.73 | 30.47 | 18.93 | 42.94 | 26.97 | 28.05 |
| 10 | 36.43 | 36.71 | 35.07 | 39.87 | 23.34 | 29.30 | 31.11 | 30.07 | 29.74 | 38.53 | 35.97 | 35.78 | 36.17 | 33.28 | 18.95 | 30.33 | 29.27 | 19.69 | 40.70 | 27.30 | 27.82 |
| 11 | 29.27 | 29.22 | 29.38 | 30.74 | 23.58 | 24.60 | 32.99 | 26.15 | 25.79 | 28.85 | 29.26 | 31.12 | 29.32 | 31.41 | 17.41 | 27.22 | 26.32 | 15.80 | 23.57 | 18.95 | 19.42 |
| 12 | 31.25 | 31.24 | 30.70 | 30.99 | 19.73 | 22.62 | 26.74 | 23.04 | 26.25 | 31.13 | 29.90 | 29.83 | 31.61 | 19.88 | 17.84 | 22.97 | 23.94 | 20.54 | 31.58 | 24.60 | 25.78 |
| 13 | 27.15 | 28.28 | 28.11 | 28.20 | 21.16 | 21.68 | 23.70 | 24.68 | 23.26 | 28.15 | 28.06 | 28.30 | 28.27 | 21.42 | 20.04 | 23.30 | 23.25 | 19.73 | 30.88 | 23.58 | 24.92 |
| 14 | 31.93 | 30.66 | 29.18 | 30.94 | 30.88 | 24.27 | 27.11 | 27.30 | 27.32 | 29.87 | 31.95 | 31.82 | 30.52 | 39.91 | 18.77 | 27.92 | 16.53 | 28.14 | 39.87 | 31.56 | 32.99 |
| 15 | 31.57 | 31.55 | 31.89 | 36.09 | 26.88 | 23.67 | 31.72 | 19.69 | 19.77 | 32.03 | 31.94 | 31.91 | 31.80 | 30.52 | 15.80 | 21.51 | 21.10 | 26.45 | 40.50 | 31.35 | 32.07 |
| Chr total | 488.55 | 481.94 | 481.08 | 494.85 | 373.81 | 386.65 | 463.00 | 417.26 | 420.72 | 487.19 | 482.50 | 485.76 | 491.13 | 423.10 | 291.28 | 422.41 | 408.09 | 291.28 | 494.85 | 463.00 | 441.14 |
| Assembly total | 504.00 | 490.06 | 489.26 | 494.85 | 412.51 | 434.23 | 474.08 | 441.56 | 441.45 | 489.14 | 482.50 | 494.28 | 491.33 | 454.96 | 303.82 | 452.84 | 437.28 | 303.82 | 504.00 | 474.08 | 458.13 |

Chr, Chromosome; **^†^** Reassembled or scaffolding based on the reference genome CDC Bethune v3.0.

**Table S5.** Pangenome gene composition and gene ontology (GO) annotation across 17 flax cultivars.

| **Cultivar** | **Category** | **Total orthogroups** | **Total gene copies** | **Duplicate gene copies** | **GO annotated copies** | **GO annotated (%)** |
| --- | --- | --- | --- | --- | --- | --- |
| CDCBethune_v3.0 | Core | 30,792 | 37,155 | 6,363 | 20,575 | 55.38 |
|  | Soft_core | 10,651 | 11,124 | 473 | 2,407 | 21.64 |
|  | Shell | 7,990 | 15,937 | 7,947 | 2,648 | 16.62 |
|  | Rare | 462 | 923 | 461 | 49 | 5.31 |
|  | Private | 1,771 | 2,013 | 242 | 9 | 0.45 |
| **Bison** | Core | 30,792 | 37,114 | 6,322 | 20,572 | 55.43 |
|  | Soft_core | 10,657 | 11,098 | 441 | 2,392 | 21.55 |
|  | Shell | 8,190 | 15,924 | 7,734 | 2,564 | 16.10 |
|  | Rare | 254 | 269 | 15 | 35 | 13.01 |
|  | Private | 720 | 726 | 6 | 4 | 0.55 |
| **Novelty** | Core | 30,792 | 37,116 | 6,324 | 20,563 | 55.40 |
|  | Soft_core | 10,663 | 11,061 | 398 | 2,415 | 21.83 |
|  | Shell | 8,126 | 15,684 | 7,558 | 2,668 | 17.01 |
|  | Rare | 242 | 251 | 9 | 36 | 14.34 |
|  | Private | 533 | 536 | 3 | 6 | 1.12 |
| T397 | Core | 30,792 | 36,857 | 6,065 | 20,392 | 55.33 |
|  | Soft_core | 9,565 | 9,937 | 372 | 1,856 | 18.68 |
|  | Shell | 6,494 | 12,554 | 6,060 | 1,603 | 12.77 |
|  | Rare | 790 | 1,711 | 921 | 91 | 5.32 |
|  | Private | 1,508 | 1,577 | 69 | 32 | 2.03 |
| K-1531 | Core | 30,792 | 37,420 | 6,628 | 20,758 | 55.47 |
|  | Soft_core | 10,589 | 11,030 | 441 | 2,383 | 21.60 |
|  | Shell | 6,220 | 6,946 | 726 | 1,289 | 18.56 |
|  | Rare | 340 | 352 | 12 | 71 | 20.17 |
|  | Private | 522 | 764 | 242 | 32 | 4.19 |
| Line3896 | Core | 30,792 | 37,408 | 6,616 | 20,638 | 55.17 |
|  | Soft_core | 10,624 | 11,050 | 426 | 2,399 | 21.71 |
|  | Shell | 6,845 | 9,167 | 2,322 | 1,619 | 17.66 |
|  | Rare | 399 | 593 | 194 | 81 | 13.66 |
|  | Private | 1,431 | 1,805 | 374 | 25 | 1.39 |
| Neiya9 | Core | 30,792 | 38,732 | 7,940 | 21,425 | 55.32 |
|  | Soft_core | 10,533 | 11,650 | 1,117 | 2,444 | 20.98 |
|  | Shell | 6,534 | 9,141 | 2,607 | 1,689 | 18.48 |
|  | Rare | 405 | 476 | 71 | 75 | 15.76 |
|  | Private | 581 | 696 | 115 | 50 | 7.18 |
| Attila | Core | 30,792 | 37,210 | 6,418 | 20,623 | 55.42 |
|  | Soft_core | 10,592 | 10,980 | 388 | 2,383 | 21.70 |
|  | Shell | 7,369 | 11,157 | 3,788 | 1,625 | 14.56 |
|  | Rare | 380 | 429 | 49 | 81 | 18.88 |
|  | Private | 665 | 729 | 64 | 26 | 3.57 |
| Marquise | Core | 30,792 | 37,197 | 6,405 | 20,623 | 55.44 |
|  | Soft_core | 10,606 | 11,065 | 459 | 2,392 | 21.62 |
|  | Shell | 7,167 | 11,066 | 3,899 | 1,711 | 15.46 |
|  | Rare | 406 | 498 | 92 | 97 | 19.48 |
|  | Private | 422 | 486 | 64 | 14 | 2.88 |
| K-3018 | Core | 30,792 | 37,173 | 6,381 | 20,601 | 55.42 |
|  | Soft_core | 10,686 | 11,116 | 430 | 2,423 | 21.80 |
|  | Shell | 8,293 | 16,262 | 7,969 | 2,808 | 17.27 |
|  | Rare | 170 | 178 | 8 | 31 | 17.42 |
|  | Private | 497 | 499 | 2 | 5 | 1.00 |
| Gaosi | Core | 30,792 | 37,084 | 6,292 | 20,544 | 55.40 |
|  | Soft_core | 10,677 | 11,048 | 371 | 2,417 | 21.88 |
|  | Shell | 8,163 | 16,337 | 8,174 | 2,864 | 17.53 |
|  | Rare | 170 | 180 | 10 | 23 | 12.78 |
|  | Private | 574 | 579 | 5 | 3 | 0.52 |
| **Laura** | Core | 30,792 | 37,077 | 6,285 | 20,548 | 55.42 |
|  | Soft_core | 10,677 | 11,116 | 439 | 2,422 | 21.79 |
|  | Shell | 8,295 | 16,456 | 8,161 | 2,838 | 17.25 |
|  | Rare | 150 | 152 | 2 | 24 | 15.79 |
|  | Private | 579 | 580 | 1 | 0 | 0.00 |
| Atlant_v2.0 | Core | 30,792 | 37,116 | 6,324 | 20,564 | 55.40 |
|  | Soft_core | 10,669 | 11,077 | 408 | 2,420 | 21.85 |
|  | Shell | 8,266 | 16,612 | 8,346 | 2,759 | 16.61 |
|  | Rare | 166 | 170 | 4 | 36 | 21.18 |
|  | Private | 813 | 822 | 9 | 3 | 0.37 |
| YY5_v2.0 | Core | 30,792 | 37,518 | 6,726 | 20,790 | 55.41 |
|  | Soft_core | 10,515 | 11,017 | 502 | 2,374 | 21.55 |
|  | Shell | 8,103 | 17,648 | 9,545 | 2,949 | 16.71 |
|  | Rare | 486 | 721 | 235 | 52 | 7.21 |
|  | Private | 1,719 | 1,754 | 35 | 9 | 0.51 |
| Heiya14 | Core | 30,792 | 37,930 | 7,138 | 21,083 | 55.58 |
|  | Soft_core | 3,118 | 3,282 | 164 | 1,640 | 49.97 |
|  | Shell | 2,158 | 2,276 | 118 | 1,040 | 45.69 |
|  | Rare | 121 | 129 | 8 | 67 | 51.94 |
|  | Private | 547 | 609 | 62 | 109 | 17.90 |
| Bolchoi | Core | 30,792 | 37,102 | 6,310 | 20,554 | 55.40 |
|  | Soft_core | 10,674 | 11,067 | 393 | 2,427 | 21.93 |
|  | Shell | 7,681 | 11,793 | 4,112 | 2,323 | 19.70 |
|  | Rare | 297 | 371 | 74 | 51 | 13.75 |
|  | Private | 484 | 561 | 77 | 22 | 3.92 |
| Ideo | Core | 30,792 | 36,762 | 5,970 | 20,351 | 55.36 |
|  | Soft_core | 9,640 | 10,044 | 404 | 1,913 | 19.05 |
|  | Shell | 7,321 | 11,276 | 3,955 | 2,276 | 20.18 |
|  | Rare | 252 | 351 | 99 | 59 | 16.81 |
|  | Private | 273 | 317 | 44 | 8 | 2.52 |

**Table S6.** Summary of morphotype-associated gene presence–absence variation (PAV) (See separate excel file)

**Table S7.** Genome-wide repeat annotation and classification across the flax pangenome, including three newly sequenced cultivars (Novelty, Bison, and Laura) and 14 previously published genome assemblies.

| **Class** | **Subclass** | **Superfamily** | **CDC Bethune v3.0^†^** | | | **Bison** | | | **Novelty** | | | | **T397** | | | | | **Attila** | | | | **K-1531** | | |
| --- | --- | --- | --- | --- | --- | --- | --- | --- | --- | --- | --- | --- | --- | --- | --- | --- | --- | --- | --- | --- | --- | --- | --- | --- |
|  |  |  | **No. of hits** | **Length (Mb)** | **%** | **No. of hits** | **Length (Mb)** | **%** | **No. of hits** | **Length (Mb)** | | **%** | | **No. of hits** | **Length (Mb)** | **%** | **No. of hits** | | **Length (Mb)** | **%** | **No. of hits** | | **Length (Mb)** | **%** |
| **Class I:Retrotransposon** | | | **242,931** | **71.69** | **14.22** | **237,036** | **70.83** | **14.45** | **228,822** | **71.13** | **14.53** | | **235,715** | | **14.53** | **14.37** | **228,968** | | **67.26** | **15.23** | **167,879** | | **63.03** | **15.29** |
|  | LTR | Copia | 26,106 | 18.00 | 3.57 | 25,833 | 17.71 | 3.61 | 25,896 | 17.63 | 3.60 | | 25,263 | | 17.36 | 3.51 | 26,058 | | 17.81 | 4.03 | 25,824 | | 17.80 | 4.32 |
|  |  | Gypsy | 102,739 | 26.11 | 5.18 | 99,406 | 25.69 | 5.24 | 95,264 | 26.10 | 5.33 | | 99,570 | | 26.23 | 5.30 | 98,614 | | 23.38 | 5.29 | 58,338 | | 19.74 | 4.79 |
|  |  | Unknown | 102,241 | 23.01 | 4.56 | 99,965 | 22.94 | 4.68 | 95,851 | 22.94 | 4.69 | | 99,647 | | 23.24 | 4.70 | 92,556 | | 21.67 | 4.91 | 72,061 | | 21.00 | 5.09 |
|  | nonLTR | LINE | 11,845 | 4.58 | 0.91 | 11,832 | 4.50 | 0.92 | 11,811 | 4.46 | 0.91 | | 11,235 | | 4.25 | 0.86 | 11,740 | | 4.40 | 1.00 | 11,656 | | 4.49 | 1.09 |
| **Class II: DNA Transposon** | | | **444,155** | **182.10** | **36.12** | **439,045** | **181.60** | **37.06** | **439,523** | **179.40** | **36.67** | | **424,799** | | **36.67** | **39.06** | **406,762** | | **143.15** | **32.43** | **373,331** | | **129.86** | **31.48** |
|  | TIR | CACTA | 75,523 | 23.81 | 4.72 | 75,937 | 23.70 | 4.84 | 73,078 | 22.54 | 4.61 | | 71,848 | | 21.37 | 4.32 | 67,187 | | 20.00 | 4.53 | 68,802 | | 20.66 | 5.01 |
|  |  | Mutator | 172,995 | 48.84 | 9.69 | 171,288 | 49.94 | 10.19 | 170,431 | 49.30 | 10.08 | | 167,319 | | 49.78 | 10.06 | 159,826 | | 43.83 | 9.93 | 133,157 | | 38.08 | 9.23 |
|  |  | PIF_Harbinger | 31,133 | 12.67 | 2.51 | 28,354 | 10.76 | 2.20 | 32,081 | 14.21 | 2.90 | | 29,688 | | 11.13 | 2.25 | 26,268 | | 8.37 | 1.90 | 22,211 | | 6.04 | 1.47 |
|  |  | Tc1_Mariner | 16,104 | 2.16 | 0.43 | 16,087 | 2.16 | 0.44 | 16,303 | 2.18 | 0.45 | | 14,422 | | 2.06 | 0.42 | 15,088 | | 2.06 | 0.47 | 13,270 | | 1.96 | 0.47 |
|  |  | hAT | 42,921 | 73.79 | 14.64 | 42,728 | 74.35 | 15.17 | 42,608 | 70.51 | 14.41 | | 40,269 | | 89.02 | 17.99 | 41,232 | | 48.96 | 11.09 | 38,580 | | 43.18 | 10.47 |
|  | nonTIR | Helitron | 105,209 | 20.81 | 4.13 | 104,381 | 20.67 | 4.22 | 104,764 | 20.65 | 4.22 | | 100,995 | | 19.88 | 4.02 | 96,905 | | 19.90 | 4.51 | 97,045 | | 19.91 | 4.83 |
|  | Other |  | 270 | 0.02 | 0.00 | 270 | 0.02 | 0.00 | 258 | 0.02 | 0.00 | | 258 | | 0.02 | 0.00 | 256 | | 0.02 | 0.00 | 266 | | 0.02 | 0.00 |
| Unknown |  |  | 374,189 | 78.69 | 15.61 | 352,639 | 66.79 | 13.63 | 351,671 | 67.98 | 13.90 | | 341,155 | | 64.64 | 13.06 | 321,501 | | 59.34 | 13.44 | 247,923 | | 47.25 | 11.46 |
| Total repeats | |  | **1,061,275** | **332.47** | **65.97** | **1,028,720** | **319.23** | **65.14** | **1,020,016** | **318.52** | **65.10** | | **65.10** | | **328.99** | **66.48** | **957,231** | | **269.75** | **61.09** | **789,133** | | **240.14** | **58.22** |
| Total non-repeats | |  |  | **171.53** | **34.03** |  | **170.83** | **34.86** |  | **170.74** | **34.90** | | **34.90** | | **165.86** | **33.52** |  | | **171.81** | **38.91** |  | | **172.29** | **41.77** |

| **Class** | **Subclass** | **Superfamily** | **K-3018** | | | **Marquise** | | | **Neiya9** | | | **Line3896** | | | **Gaosi** |  |  |
| --- | --- | --- | --- | --- | --- | --- | --- | --- | --- | --- | --- | --- | --- | --- | --- | --- | --- |
|  |  |  | **No. of hits** | **Length (Mb)** | **%** | **No. of hits** | **Length (Mb)** | **%** | **No. of hits** | **Length (Mb)** | **%** | **No. of hits** | **Length (Mb)** | **%** | **No. of hits** | **Length (Mb)** | **%** |
| **Class I:Retrotransposon** | | | **252,061** | **72.85** | **14.90** | **221,442** | **69.75** | **15.80** | **226,790** | **75.12** | **15.85** | **207,596** | **66.91** | **15.41** | **238,931** | **71.36** | **14.79** |
|  | LTR | Copia | 25,892 | 17.85 | 3.65 | 26,093 | 17.92 | 4.06 | 27,410 | 18.66 | 3.94 | 25,757 | 17.61 | 4.05 | 25,983 | 17.86 | 3.70 |
|  |  | Gypsy | 106,642 | 26.78 | 5.48 | 99,532 | 26.11 | 5.92 | 93,876 | 26.61 | 5.61 | 84,605 | 23.10 | 5.32 | 99,728 | 26.12 | 5.41 |
|  |  | Unknown | 107,658 | 23.72 | 4.85 | 84,081 | 21.30 | 4.82 | 93,059 | 24.85 | 5.24 | 85,661 | 21.75 | 5.01 | 101,430 | 22.94 | 4.76 |
|  | nonLTR | LINE | 11,869 | 4.50 | 0.92 | 11,736 | 4.42 | 1.00 | 12,445 | 5.00 | 1.06 | 11,573 | 4.45 | 1.03 | 11,790 | 4.43 | 0.92 |
| **Class II: DNA Transposon** | | | **442,158** | **179.03** | **36.61** | **405,057** | **143.42** | **32.49** | **442,865** | **149.62** | **31.55** | **406,191** | **134.79** | **31.04** | **437,046** | **174.67** | **36.21** |
|  | TIR | CACTA | 71,023 | 21.52 | 4.40 | 67,254 | 19.88 | 4.50 | 79,595 | 25.18 | 5.31 | 72,198 | 22.66 | 5.22 | 70,319 | 21.09 | 4.37 |
|  |  | Mutator | 177,588 | 50.12 | 10.25 | 160,022 | 43.61 | 9.88 | 168,093 | 46.16 | 9.74 | 149,271 | 41.10 | 9.47 | 173,028 | 50.50 | 10.47 |
|  |  | PIF_Harbinger | 30,873 | 10.41 | 2.13 | 26,136 | 8.03 | 1.82 | 30,598 | 8.67 | 1.83 | 26,507 | 7.43 | 1.71 | 31,347 | 12.33 | 2.56 |
|  |  | Tc1_Mariner | 16,241 | 2.18 | 0.45 | 13,055 | 1.92 | 0.44 | 13,740 | 2.01 | 0.42 | 15,620 | 2.11 | 0.49 | 16,114 | 2.16 | 0.45 |
|  |  | hAT | 42,527 | 74.29 | 15.19 | 39,486 | 49.57 | 11.23 | 42,689 | 45.53 | 9.60 | 41,935 | 41.09 | 9.46 | 42,688 | 68.16 | 14.13 |
|  | nonTIR | Helitron | 103,654 | 20.48 | 4.19 | 98,832 | 20.39 | 4.62 | 107,892 | 22.05 | 4.65 | 100,399 | 20.37 | 4.69 | 103,303 | 20.40 | 4.23 |
|  | Other |  | 252 | 0.02 | 0.00 | 272 | 0.02 | 0.00 | 258 | 0.02 | 0.00 | 261 | 0.02 | 0.00 | 247 | 0.02 | 0.00 |
| Unknown |  |  | 365,076 | 66.27 | 13.55 | 322,198 | 56.42 | 12.78 | 347,807 | 71.52 | 15.09 | 303,631 | 58.13 | 13.39 | 351,670 | 65.56 | 13.59 |
| Total repeats | |  | **1,059,295** | **318.14** | **65.04** | **948,697** | **269.60** | **61.07** | **1,017,462** | **296.26** | **62.49** | **917,418** | **259.82** | **59.84** | **1,027,647** | **311.58** | **64.58** |
| Total non-repeats | |  |  | **170.99** | **34.96** |  | **171.85** |  |  | **177.82** | **37.51** |  | **174.41** | **40.17** |  | **484.50** | **35.69** |

| **Class** | **Subclass** | **Superfamily** | **Laura** | | | **Atlant v2.0** | | | **YY5 v2.0** | | | **Bolchoi** | | | **Ideo** | | | **Heiya14** | | |
| --- | --- | --- | --- | --- | --- | --- | --- | --- | --- | --- | --- | --- | --- | --- | --- | --- | --- | --- | --- | --- |
|  |  |  | **No. of hits** | **Length (Mb)** | **%** | **No. of hits** | **Length (Mb)** | **%** | **No. of hits** | **Length (Mb)** | **%** | **No. of hits** | **Length (Mb)** | **%** | **No. of hits** | **Length (Mb)** | **%** | **No. of hits** | **Length (Mb)** | % |
| Calss I:Retrotransposon | | | **237,219** | **71.91** | **14.54** | **235,567** | **71.58** | **14.57** | **244,732** | **72.87** | **16.02** | **219,174** | **69.23** | **15.29** | **231,402** | **70.20** | **16.04** | **102,238** | **47.70** | **15.70** |
|  | LTR | Copia | 25,891 | 17.76 | 3.59 | 25,880 | 17.78 | 3.62 | 25,821 | 17.77 | 3.91 | 25,979 | 17.77 | 3.92 | 25,557 | 17.55 | 4.01 | 24,702 | 15.88 | 5.23 |
|  |  | Gypsy | 99,848 | 26.51 | 5.36 | 98,868 | 26.47 | 5.39 | 101,865 | 26.62 | 5.85 | 95,895 | 25.28 | 5.58 | 100,715 | 26.17 | 5.98 | 23,354 | 12.07 | 3.97 |
|  |  | Unknown | 99,612 | 23.13 | 4.68 | 99,028 | 22.87 | 4.65 | 105,184 | 24.02 | 5.28 | 85,609 | 21.76 | 4.81 | 93,559 | 22.10 | 5.05 | 42,949 | 15.33 | 5.05 |
|  | nonLTR | LINE | 11,868 | 4.51 | 0.91 | 11,791 | 4.45 | 0.91 | 11,862 | 4.47 | 0.98 | 11,691 | 4.42 | 0.98 | 11,571 | 4.38 | 1.00 | 11,233 | 4.42 | 1.45 |
| Class II: DNA Transposon | | | **441,959** | **180.44** | **36.49** | **440,405** | **181.76** | **37.00** | **429,636** | **141.59** | **31.13** | **413,417** | **143.80** | **31.75** | **404,679** | **136.39** | **31.18** | **262,319** | **59.31** | **19.53** |
|  | TIR | CACTA | 72,747 | 22.06 | 4.46 | 74,567 | 22.39 | 4.56 | 71,606 | 20.50 | 4.51 | 68,838 | 21.06 | 4.65 | 63,764 | 19.72 | 4.51 | 54,095 | 12.59 | 4.14 |
|  |  | Mutator | 172,962 | 50.31 | 10.18 | 173,482 | 51.98 | 10.58 | 173,058 | 52.04 | 11.44 | 160,594 | 45.78 | 10.11 | 161,061 | 44.01 | 10.06 | 79,019 | 17.67 | 5.82 |
|  |  | PIF_Harbinger | 32,226 | 14.11 | 2.85 | 29,070 | 11.82 | 2.41 | 31,743 | 14.83 | 3.26 | 25,521 | 7.84 | 1.73 | 26,187 | 7.62 | 1.74 | 11,481 | 2.37 | 0.78 |
|  |  | Tc1_Mariner | 16,190 | 2.17 | 0.44 | 16,180 | 2.16 | 0.44 | 12,454 | 1.90 | 0.42 | 15,575 | 2.06 | 0.45 | 15,042 | 2.00 | 0.46 | 9,231 | 1.69 | 0.56 |
|  |  | hAT | 42,785 | 71.08 | 14.38 | 42,816 | 72.92 | 14.84 | 37,841 | 31.78 | 6.99 | 42,076 | 46.85 | 10.35 | 41,702 | 43.17 | 9.87 | 31,137 | 7.40 | 2.44 |
|  | nonTIR | Helitron | 104,783 | 20.68 | 4.18 | 104,018 | 20.47 | 4.17 | 102,666 | 20.51 | 4.51 | 100,556 | 20.20 | 4.46 | 96,656 | 19.84 | 4.54 | 77,093 | 17.57 | 5.78 |
|  | Other |  | 266 | 0.02 | 0.00 | 272 | 0.02 | 0.00 | 268 | 0.02 | 0.00 | 257 | 0.02 | 0.00 | 267 | 0.02 | 0.00 | 263 | 0.03 | 0.01 |
| Unknown |  |  | 356,539 | 71.28 | 14.42 | 352,254 | 66.55 | 13.55 | 351,960 | 67.81 | 14.91 | 324,117 | 65.22 | 14.40 | 336,238 | 63.34 | 14.49 | 116,731 | 24.51 | 8.07 |
| Total repeats | |  | **1,035,717** | **323.64** | **65.48** | **1,028,226** | **319.89** | **65.11** | **1,026,328** | **282.28** | **62.04** | **956,708** | **278.25** | **61.45** | **972,319** | **269.93** | **61.73** | **481,288** | **131.52** | **43.29** |
| Total non-repeats | |  |  | **170.64** | **34.52** |  | **171.44** | **34.89** |  | **172.68** | **37.96** |  | **174.59** | **38.55** |  | **167.35** | **38.27** |  | **172.3** | **56.71** |

**Table S8.** Chromosome-level summary statistics of population differentiation and nucleotidediversity ($\pi$) in fiber and linseed flax.

| **Chr** | **No. windows** | **Mean window SNP count** | **SNP count *CV*** | **Mean weighted** $\boldsymbol{F}_{\boldsymbol{ST}}$ | **Weight** $\boldsymbol{F}_{\boldsymbol{ST}}$ ***CV*** | **Mean linseed** $\boldsymbol{\pi}$ | **Linseed** $\boldsymbol{\pi}$ ***CV*** | **Mean fiber flax** $\boldsymbol{\pi}$ | **Fiber flax** $\boldsymbol{\pi}$ ***CV*** | **Mean** $\boldsymbol{\pi}_{\boldsymbol{linseed}}\mathbf{/}\boldsymbol{\pi}_{\boldsymbol{fiber flax}}$ | $\boldsymbol{\pi}_{\boldsymbol{linseed}}\mathbf{/}\boldsymbol{\pi}_{\boldsymbol{fiber} \boldsymbol{flax}}$ ***CV*** | **Mean** $\boldsymbol{\pi}_{\boldsymbol{fiber flax}}\mathbf{/}\boldsymbol{\pi}_{\boldsymbol{linseed}}$ | $\boldsymbol{\pi}_{\boldsymbol{fiber} \boldsymbol{flax}}\mathbf{/}\boldsymbol{\pi}_{\boldsymbol{linseed}}$***CV*** | **Mean log₂(**$\boldsymbol{\pi}_{\boldsymbol{linseed}}\mathbf{/}\boldsymbol{\pi}_{\boldsymbol{fiber flax}}$**)** | **log₂(**$\boldsymbol{\pi}_{\boldsymbol{linseed}}\mathbf{/}\boldsymbol{\pi}_{\boldsymbol{fiber} \boldsymbol{flax}}$**) *CV*** |
| --- | --- | --- | --- | --- | --- | --- | --- | --- | --- | --- | --- | --- | --- | --- | --- |
| Chr1 | 2,996 | 466 | 0.88 | 0.1545 | 0.79 | 0.0015 | 0.95 | 0.0009 | 1.11 | 2.63 | 2.79 | 0.73 | 0.99 | 0.70 | 1.38 |
| Chr2 | 3,002 | 456 | 1.17 | 0.1269 | 0.55 | 0.0014 | 1.23 | 0.0008 | 1.38 | 1.89 | 1.18 | 0.73 | 0.68 | 0.65 | 1.18 |
| Chr3 | 2,939 | 401 | 0.99 | 0.0842 | 0.73 | 0.0012 | 1.09 | 0.0008 | 1.28 | 2.14 | 2.39 | 0.89 | 0.54 | 0.42 | 2.39 |
| Chr4 | 2,407 | 259 | 1.20 | 0.0884 | 0.59 | 0.0008 | 1.31 | 0.0006 | 1.41 | 1.31 | 0.64 | 0.93 | 0.51 | 0.24 | 2.62 |
| Chr5 | 2,354 | 540 | 0.89 | 0.0965 | 0.58 | 0.0016 | 0.96 | 0.0011 | 1.07 | 1.94 | 1.31 | 0.86 | 0.86 | 0.52 | 1.86 |
| Chr6 | 1,938 | 552 | 0.74 | 0.0950 | 0.67 | 0.0017 | 0.80 | 0.0012 | 0.88 | 1.48 | 0.43 | 0.82 | 0.83 | 0.45 | 1.34 |
| Chr7 | 2,081 | 500 | 1.01 | 0.1289 | 0.51 | 0.0016 | 1.09 | 0.0010 | 1.14 | 1.75 | 0.60 | 0.67 | 0.35 | 0.67 | 0.85 |
| Chr8 | 3,446 | 203 | 1.73 | 0.0739 | 0.71 | 0.0006 | 1.85 | 0.0005 | 1.90 | 1.15 | 0.46 | 1.03 | 0.62 | 0.09 | 6.41 |
| Chr9 | 2,781 | 475 | 1.00 | 0.1370 | 0.55 | 0.0015 | 1.06 | 0.0010 | 1.17 | 1.74 | 0.94 | 0.90 | 0.84 | 0.45 | 2.13 |
| Chr10 | 2,595 | 388 | 1.06 | 0.1013 | 0.54 | 0.0012 | 1.17 | 0.0008 | 1.24 | 2.07 | 1.54 | 0.78 | 0.84 | 0.65 | 1.41 |
| Chr11 | 2,295 | 364 | 1.04 | 0.1155 | 0.48 | 0.0012 | 1.11 | 0.0007 | 1.22 | 2.10 | 3.37 | 0.76 | 1.14 | 0.66 | 1.25 |
| Chr12 | 2,487 | 518 | 0.92 | 0.1262 | 0.53 | 0.0016 | 1.02 | 0.0009 | 1.13 | 1.78 | 0.82 | 0.88 | 0.81 | 0.52 | 1.85 |
| Chr13 | 2,069 | 759 | 0.53 | 0.1112 | 0.49 | 0.0024 | 0.56 | 0.0015 | 0.71 | 2.15 | 0.83 | 0.60 | 0.39 | 0.87 | 0.82 |
| Chr14 | 2,408 | 472 | 0.95 | 0.1581 | 0.62 | 0.0015 | 0.99 | 0.0010 | 1.16 | 1.99 | 1.95 | 0.70 | 0.56 | 0.68 | 1.12 |
| Chr15 | 2,023 | 581 | 0.69 | 0.1656 | 0.59 | 0.0019 | 0.70 | 0.0010 | 0.95 | 3.00 | 0.91 | 0.50 | 0.56 | 1.25 | 0.74 |

Chr: chromosome; *CV*, coefficient of variation, defined as standard deviation divided by mean.

**Table S9.** Summary of quantitative trait nucleotides (QTNs) identified in the whole diversity population and their associated candidate genes.

| **Trait group** | **Trait ID** | **Trait full name** | **Total QTNs** | **Min *R*^2^** | **Max *R*^2^** | **Mean *R*^2^** | **Mean MAF** | **QTNs in gene** | **QTNs within 5kb** | **QTNs within 50kb** | **Genes in gene** | **Genes within 5kb** | **Genes within 50kb** |
| --- | --- | --- | --- | --- | --- | --- | --- | --- | --- | --- | --- | --- | --- |
| Architecture | PLH | Plant height (cm) | 92 | 0.23 | 5.36 | 1.40 | 0.24 | 20 | 84 | 92 | 20 | 238 | 2,032 |
|  | BSC | Branching score | 71 | 0.00 | 9.86 | 2.33 | 0.28 | 15 | 67 | 71 | 15 | 180 | 1,517 |
|  | LOD | Lodging score | 61 | 0.00 | 15.87 | 2.96 | 0.24 | 15 | 56 | 61 | 13 | 152 | 1,212 |
| Fiber | LIG | Lignin content (%) | 181 | 0.00 | 8.63 | 0.39 | 0.21 | 43 | 165 | 181 | 43 | 452 | 3,449 |
|  | SHI | Shive content (%) | 115 | 0.20 | 10.64 | 1.92 | 0.24 | 26 | 100 | 115 | 25 | 264 | 2,273 |
|  | FIB | Fiber content (%) | 111 | 0.24 | 12.52 | 2.00 | 0.25 | 25 | 92 | 111 | 23 | 251 | 2,160 |
|  | CEW | Cell wall content (%) | 92 | 0.00 | 10.18 | 2.44 | 0.24 | 24 | 88 | 92 | 24 | 231 | 1,978 |
|  | CEL | Cellulose content(%) | 85 | 0.48 | 9.14 | 2.25 | 0.28 | 22 | 84 | 85 | 21 | 219 | 1,816 |
|  | STR | Straw weight (g) | 74 | 0.22 | 9.31 | 1.93 | 0.24 | 24 | 69 | 73 | 24 | 190 | 1,516 |
| Oil | OIL | Oil content (%) | 132 | 0.31 | 11.52 | 1.60 | 0.24 | 34 | 125 | 132 | 34 | 356 | 2,724 |
|  | LIO | Linoleic acid content (%) | 129 | 0.19 | 10.12 | 2.17 | 0.16 | 22 | 117 | 129 | 22 | 318 | 2,447 |
|  | PRO | Protein content (%) | 127 | 0.03 | 6.57 | 1.68 | 0.24 | 32 | 120 | 127 | 30 | 314 | 2,571 |
|  | IOD | Iodine value | 126 | 0.36 | 6.37 | 1.94 | 0.23 | 28 | 114 | 124 | 26 | 283 | 2,386 |
|  | LIN | Linolenic acid content (%) | 121 | 0.16 | 7.22 | 2.17 | 0.19 | 26 | 112 | 119 | 25 | 298 | 2,369 |
| Yield | TSW | Thousand-seed weight (g) | 115 | 0.00 | 7.63 | 1.19 | 0.23 | 33 | 105 | 115 | 33 | 296 | 2,408 |
|  | YLD | Seed yield (t/ha) | 80 | 0.00 | 11.59 | 1.00 | 0.22 | 16 | 70 | 80 | 16 | 171 | 1,600 |

Gene counts are cumulative across distance thresholds (i.e., genes within 5 kb include those within gene bodies, and genes within 50 kb include all genes within 5 kb).

For the linseed population, in addition to the full panel, GWAS was also performed using ten random subsets of 85 accessions drawn from the full set of 293 linseed accessions. The reported QTN, *R*², and minor allele frequency (MAF) values represent the mean across these random subsets.

**Table S10.** Summary statistics of morphotype-enriched genomic blocks identified through integrated analysis of population genetic signals and GWAS-derived QTNs.

| **Block type** | **No. blocks** | **Total length (Mb)** | **No. genes** | **No. fiber QTNs** | **No. linseed QTNs** | **Mean block size (Mb)** | **Mean No. genes** | **Mean No. fiber QTNs** | **Mean linseed QTNs** | **Mean Fst** | **Mean** $\boldsymbol{\pi}_{\boldsymbol{fiber flax}}\mathbf{/}\boldsymbol{\pi}_{\boldsymbol{linseed}}$ | **Mean** $\boldsymbol{\pi}_{\boldsymbol{linseed}}\mathbf{/}\boldsymbol{\pi}_{\boldsymbol{fiber flax}}$ |
| --- | --- | --- | --- | --- | --- | --- | --- | --- | --- | --- | --- | --- |
| Fiber enriched | 87 | 33.85 | 6,135 | 6 | 63 | 0.39 | 71 | 0.07 | 0.72 | 0.18 | 0.46 | 2.99 |
| Linseed enriched | 46 | 13.78 | 2,331 | 5 | 14 | 0.30 | 51 | 0.11 | 0.30 | 0.17 | 0.93 | 1.26 |
| Mixed direction | 18 | 9.73 | 1,518 | 2 | 16 | 0.54 | 84 | 0.11 | 0.89 | 0.15 | 0.74 | 2.54 |
| $F_{ST}$ only | 88 | 16.03 | 2,715 | 1 | 23 | 0.18 | 31 | 0.01 | 0.26 | 0.18 | 0.65 | 1.65 |
| $\pi$ only for fiber flax | 128 | 24.43 | 4,218 | 5 | 27 | 0.19 | 33 | 0.04 | 0.21 | 0.10 | 0.48 | 3.08 |
| $\pi$ only for linseed | 204 | 43.65 | 5,496 | 15 | 57 | 0.21 | 27 | 0.07 | 0.28 | 0.08 | 1.04 | 1.06 |
| $\pi$ only for mixed | 19 | 8.35 | 1,497 | 2 | 5 | 0.44 | 79 | 0.11 | 0.26 | 0.09 | 0.71 | 3.29 |
| Mean |  |  |  |  |  | 0.32 | 54 | 0.07 | 0.42 | 0.14 | 0.72 | 2.27 |

**Table S11.** Detailed summary of morphotype-enriched genomic blocks (See separate excel file)

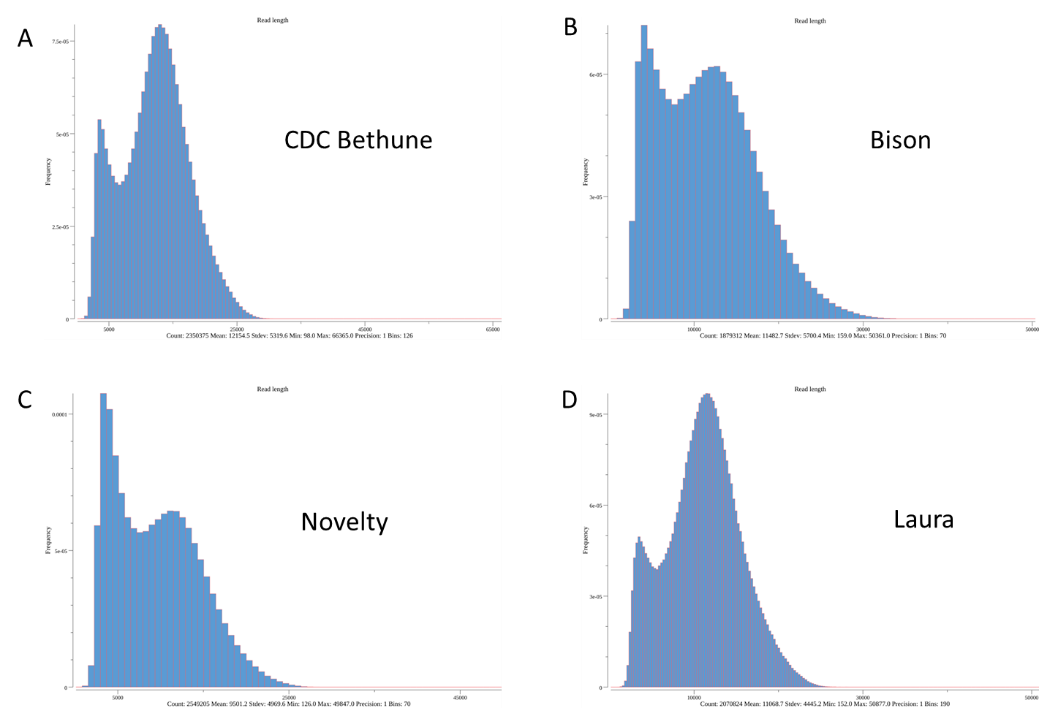

**Fig. S1**. Distribution of PacBio HiFi reads generated from four flax genomes. CDC Bethune v3.0 (You et al. 2026).

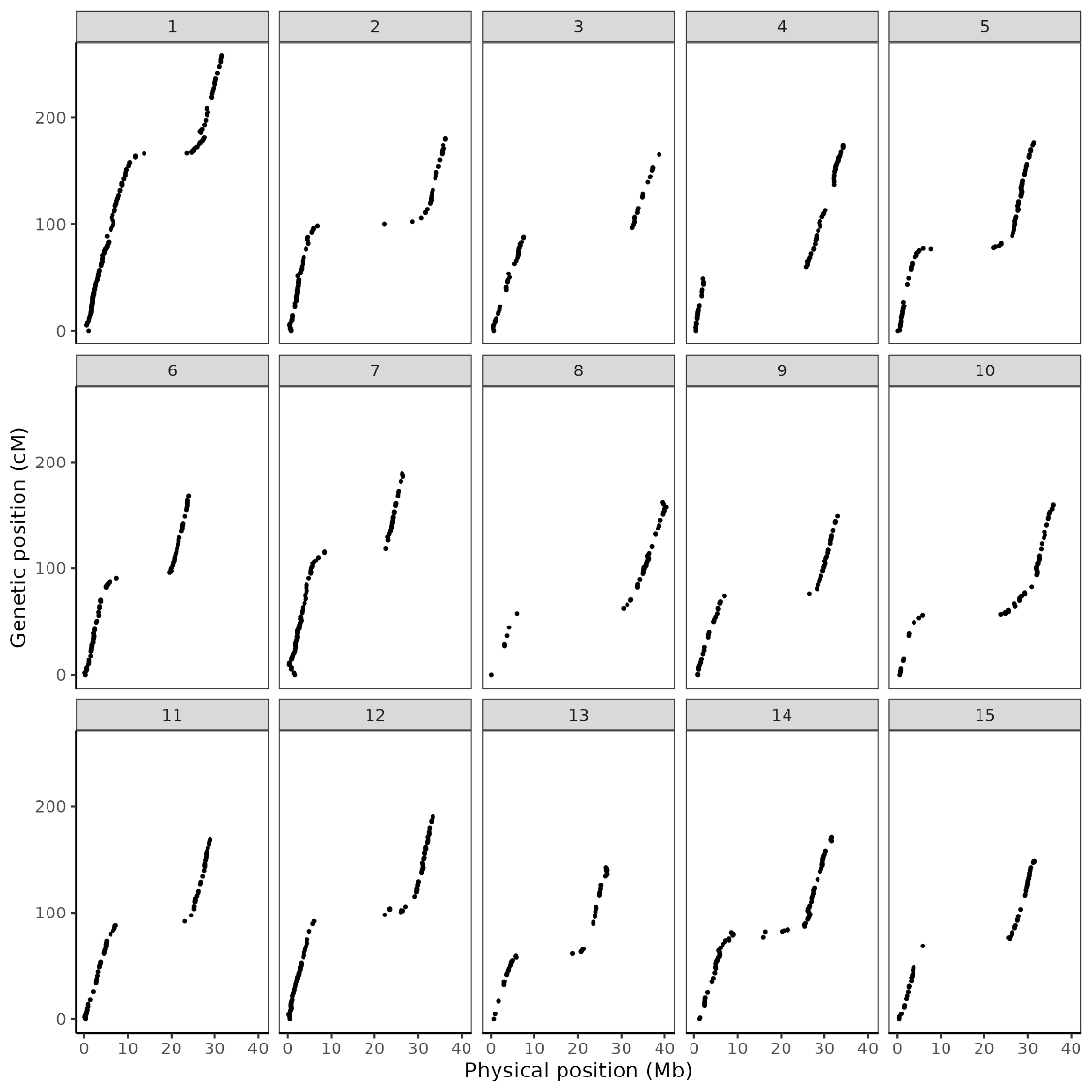

**Fig. S2.** Colinear relationship between genetic and physical distances based on the Bison x Novelty recombinant inbreeding line (RIL) population genetic map.

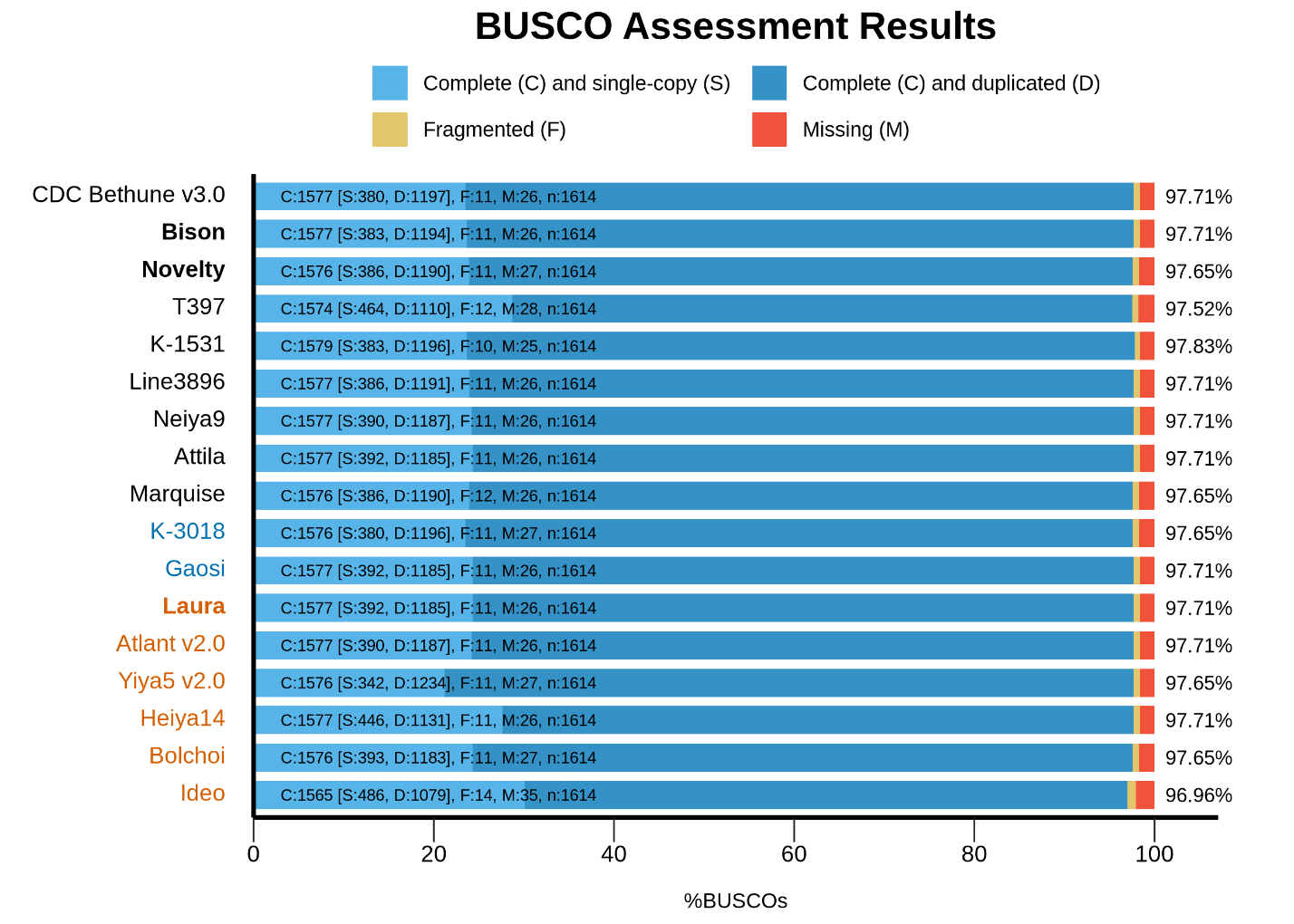

**Fig. S3.** BUSCO assessment of genome completeness for three flax cultivars newly sequenced in this study (bold) and 14 previously published genome assemblies, representing nine linseed (black), two dual (blue) and six fiber flax (orange) cultivars. The total BUSCO scores ranged from 97.0% to 97.8% and are presented at right side of each bar.

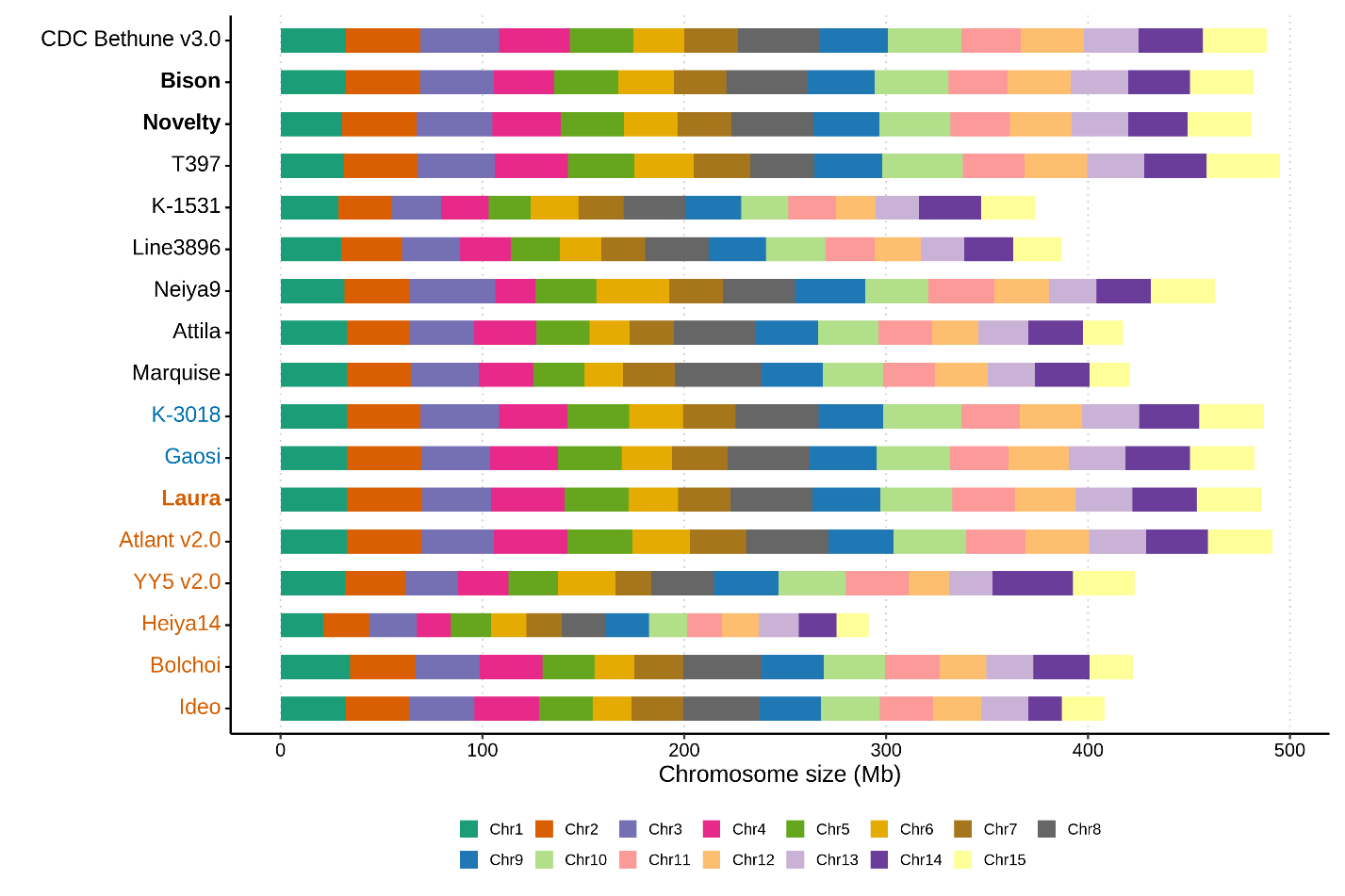

**Fig. S4.** Chromosome length distribution across 17 flax assemblies, including three flax cultivars newly sequenced in this study (bold) and 14 previously published genome assemblies, representing nine linseed (black), two dual (blue) and six fiber flax (orange) cultivars.

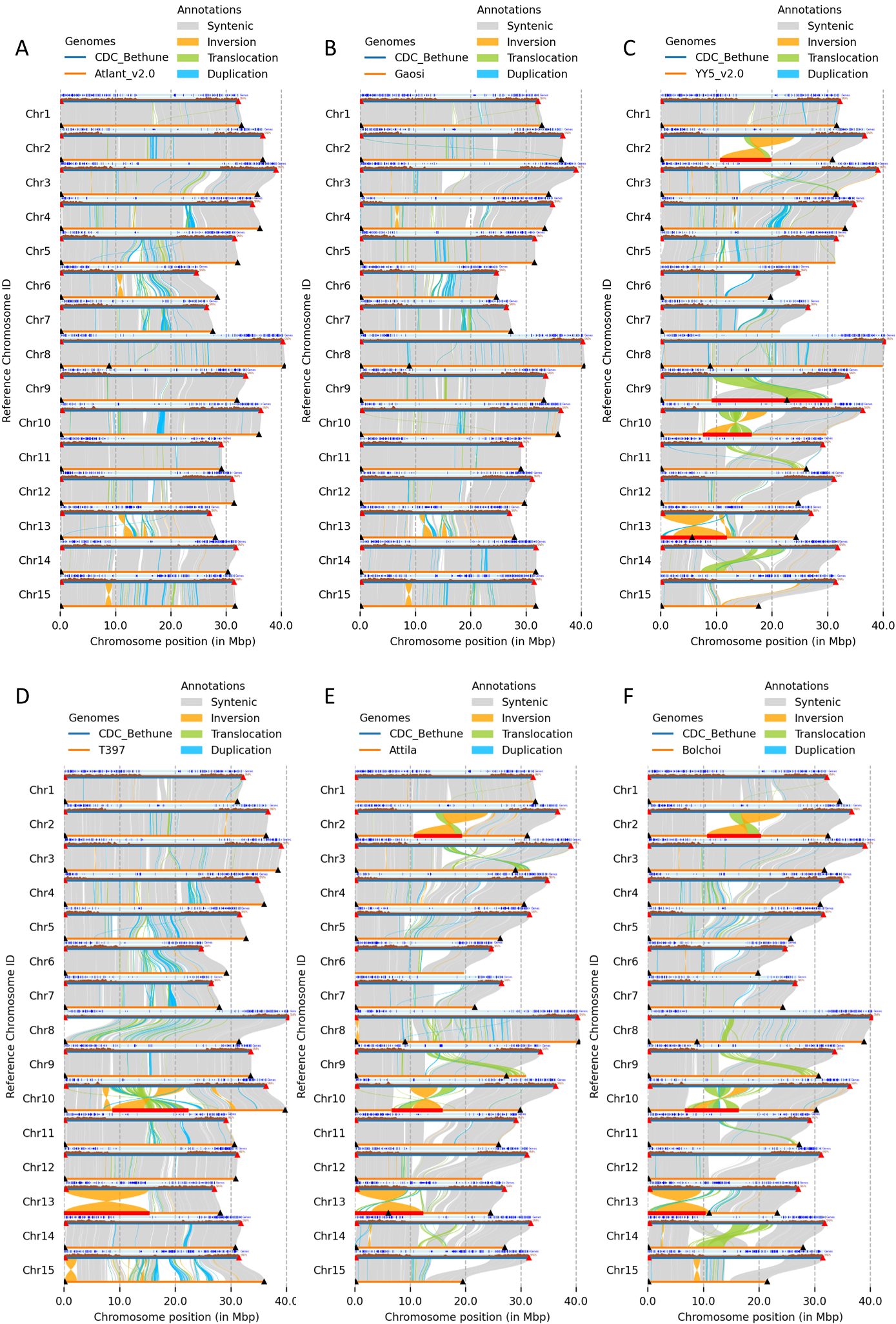

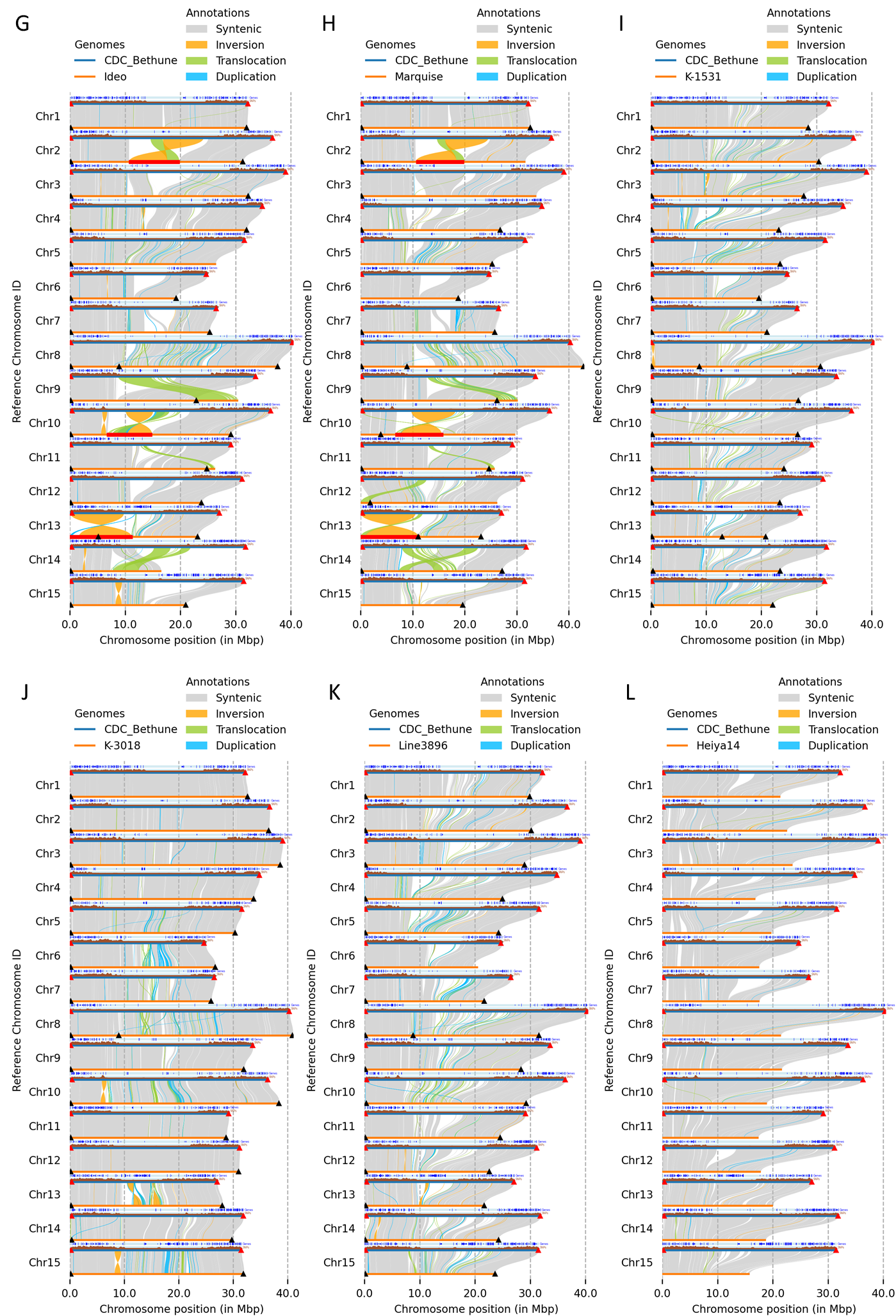

**Fig. S5**. Genome-wide structural variation between CDC Bethune v3.0 and 12 previously published flax cultivar assemblies. **A:** Russian fiber cultivar Atlant v2.0, **B:** Chinese dual-purpose cultivar Gaosi, **C:** Chinese fiber cultivar YY5 v2.0, **D:** Indian linseed cultivar T397, **E:** linseed cultivar Attila, **F:** fiber cultivar Bolchoi, **G**: cultivar Ideo, **H:** cultivar Marquise, **I:** cultivar K-1531, **J:** linseed cultivar K-3018, **K:** linseed cultivar Line3896, and **L:** fiber cultivar Heiya14. Syntenic regions, inversions, translocations, and duplications are displayed across chromosomes 1–15. Triangles indicate the positions of telomeric repeat arrays at chromosome termini. Potential assembly artifacts are highlighted with red lines. Chromosome identifiers were standardized according to the CDC Bethune v3.0 reference genome. Assemblies of K-1531 and K-3018 were generated using RagTag with CDC Bethune v3.0 as the reference genome, and assemblies of K-3018, Line3896, and Heiya14 were regenerated using CDC Bethune v3.0 as the reference genome.

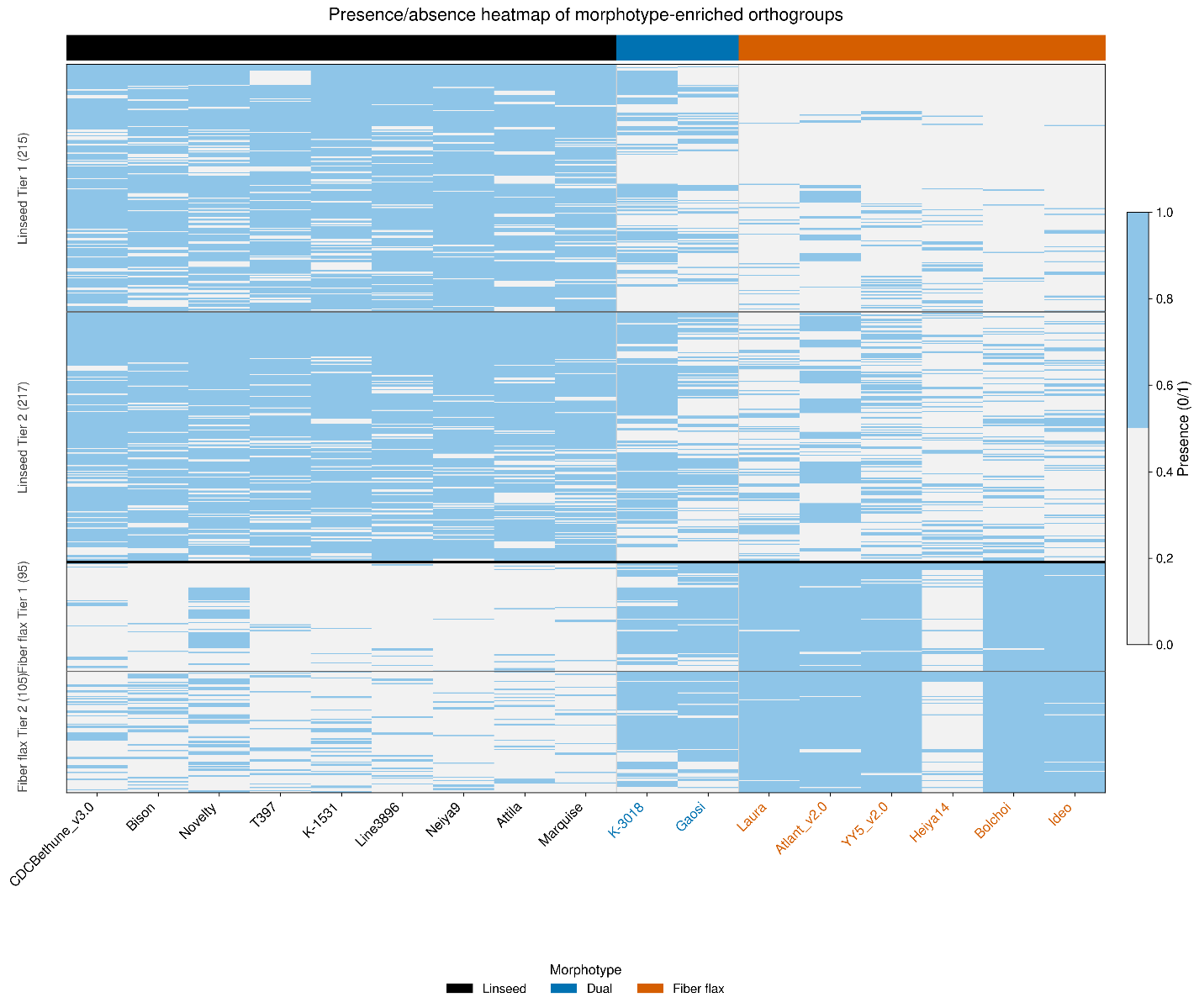

**Fig. S6.** Presence–absence variation (PAV) heatmap of morphotype-enriched orthogroups identified from flax pangenome analysis. Orthogroups were classified into Linseed Tier 1 (215), Linseed Tier 2 (217), Fiber flax Tier 1 (95), and Fiber flax Tier 2 (105) categories according to predefined morphotype-enrichment criteria. Enrichment analysis was performed using nine linseed and six fiber flax cultivars, excluding the two dual-purpose cultivars (K-3018 and Gaosi). The heatmap includes all 17 cultivated flax genomes to visualize the distribution of candidate orthogroups across morphotypes. Blue cells indicate presence and gray cells indicate absence of orthogroups. Cultivar labels are colored according to morphotype: linseed (black), dual-purpose (blue), and fiber flax (orange).

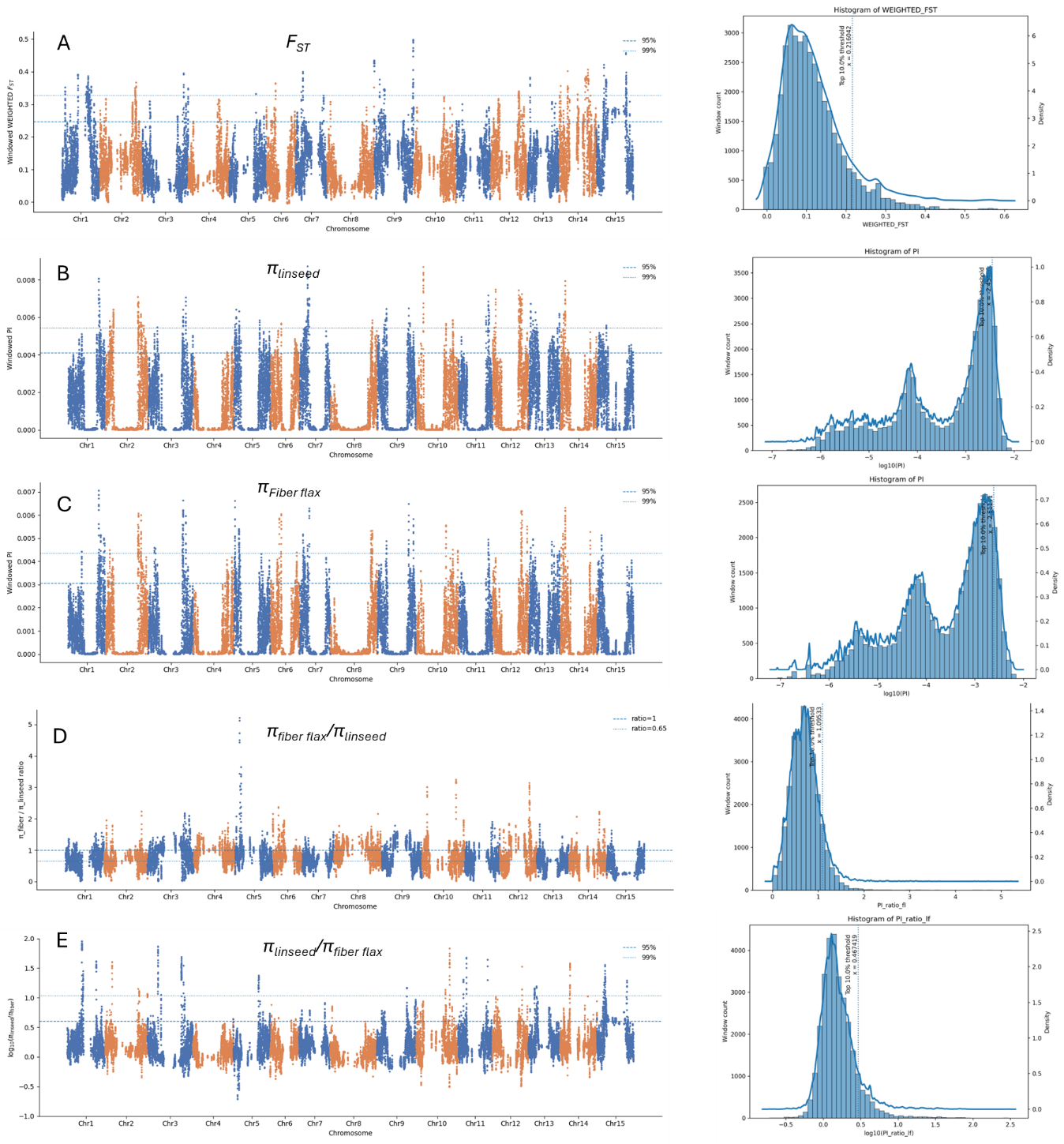

**Fig. S7.** Genome-wide nucleotide diversity and genetic differentiation between fiber and linseed flax. **A:** Genome-wide genetic differentiation ($F_{ST}$) between fiber and linseed flax, showing heterogeneous differentiation across chromosomes with several peaks exceeding the 95% and 99% thresholds, indicative of candidate regions under morphotype-specific selection. **B–C:** Genome-wide distribution of nucleotide diversity (π) in linseed (A) and fiber (B) flax across chromosomes 1–15, calculated in sliding windows. Horizontal dashed lines indicate the 95% and 99% percentile thresholds. **D:** Genome-wide ratio of nucleotide diversity between morphotypes ($\pi_{fiber flax}/\pi_{linseed}$). Most windows cluster around ~0.6–0.7, indicating a consistent reduction in diversity in fiber flax, with occasional deviations suggesting localized selection. **E:** Reciprocal diversity ratio ($\pi_{linseed}/\pi_{fiber flax}$), highlighting regions with elevated diversity in linseed relative to fiber.
